# Melt Electrowriting of Scaffolds with a Porosity Gradient to Mimic the Matrix Structure of the Human Trabecular Meshwork

**DOI:** 10.1101/2022.05.28.476655

**Authors:** Małgorzata K. Włodarczyk-Biegun, Maria Villiou, Marcus Koch, Christina Muth, Peixi Wang, Jenna Ott, Aranzazu del Campo

## Abstract

The permeability of the Human Trabecular Meshwork (HTM) regulates eye pressure via a porosity gradient across its thickness modulated by stacked layers of matrix fibrils and cells. Changes in HTM porosity are associated with increases in intraocular pressure and the progress of diseases like glaucoma. Engineered HTMs could help to understand the structure-function relation in natural tissues, and lead to new regenerative solutions. Here, melt electrowriting (MEW) is explored as a biofabrication technique to produce fibrillar, porous scaffolds that mimic the multilayer, gradient structure of native HTM. Poly(caprolactone) constructs with a height of 125-500 μm and fiber diameters of 10-12 μm are printed. Scaffolds with a tensile modulus between 5.6 and 13 MPa, and a static compression modulus in the range of 6-360 kPa are obtained by varying the scaffolds design, i.e., density and orientation of the fibers and number of stacked layers. Primary HTM cells attach to the scaffolds, proliferate, and form a confluent layer within 8-14 days, depending on the scaffold design. High cell viability and cell morphology close to that in the native tissue are observed. The present work demonstrates the utility of MEW to reconstruct complex morphological features of natural tissues.

## 1. Introduction

Natural tissues and organs have multilayered structures with varying spatial gradients in morphology and/or composition that result in unique properties and functions [1, 2]. An example is the permeability of the Human Trabecular Meshwork (HTM), which is achieved by a distinct porosity gradient across its thickness that regulates internal ocular pressure [3]. The HTM is composed of three distinct zones that decrease in pore size: the uveal region (UVM) with pore sizes varying from 70-100 um^2^ [4], the corneoscleral region (CTM) with pore sizes of 30 um^2^, and the juxtacanalicular region (JCT) with pore sizes of 4-7 um^2^ [5, 6]. The pores are formed by the cells and the matrix organized in lamellar beams with a thickness of 5-12 µm [7, 8]. The HTM has a reported full thickness of 70 – 130 µm [9], with CTM as the main compartment, the UVM region encompassing 40.6±10.0 µm [4], and the JCT region encompassing 2-20 µm [4, 7]. However, the HTM is a triangle in a cross-section, and in the thickest region can reach over 600 µm [10]. The resistance of HTM to fluid outflow increases across its thickness in the direction of the eye surface as the porosity decreases. Both aging and certain diseases can decrease the overall porosity in the HTM, resulting in poor regulation of the aqueous humor generation and drainage, and as a consequence, increased intraocular pressure. Thus, changes in HTM porosity are a primary risk factor for glaucoma and, eventually, loss of eyesight [5, 7]. Such changes in the porosity are a consequence of alterations in the morphology and the mechanical properties of the cell-matrix layers in the HTM. Studies have shown that the elastic modulus of healthy HTM, measured locally using AFM, is ∼4 kPa, whereas in glaucomatous HTM, this value increases [11-15].

In vitro studies showed that the morphology and protein expression patterns of HTM cells cultured on stiff scaffolds resembled that of HTM cells in glaucomatous tissue [16]. A reconstruction of the HTM structure could help to understand how structure and function are correlated in the natural tissue, and eventually lead to regenerative solutions to associated diseases [17]. HTM cultures in microfabricated membranes of SU-8 photoresist with pores of 12-μm size were reported [8, 18, 19]. The cells developed an HTM phenotype in terms of morphology, expression of HTM cell-specific markers, and ECM secretion. The proposed model mimicked in vivo outflow physiology: it was responsive to latrunculin-B in a dose-dependent manner [8], and a pathological state with increased ECM accumulation and decreased tissue permeability could be induced by treatment with steroids [19] or with the fibrotic agent TGFβ-2 [18], and counteracted by a ROCK inhibitor [18, 19]. In other studies, the model based on fibrillar hydrogels of collagen/elastin-like peptides was used [13], and the pathological state was successfully induced by dexamethasone, and attenuated by ROCK inhibitor, as revealed by the increased contractility, fibronectin deposition, and hydrogel stiffening. Collagen and collagen/chondroitin sulfate scaffolds obtained via freeze-drying were also used to build in vitro HTM models. Native HTM cells after 14 days of culture were viable and proliferated on the surface, invaded the scaffolds, and stretched along the collagen fibers [20]. On collagen/hyaluronic acid scaffolds with different pore sizes and connectivity obtained via freeze-drying [21], HTM cells proliferated more in larger pores. Fibronectin expression was upregulated with increasing GAGs incorporation, and the morphology of secreted fibronectin was affected by the pore architecture. A 3D culture in Matrigel [22, 23] revealed the ability of HTM cells to adapt to chronic oxidative stress, and this was more efficient in dynamic cell culture conditions [22, 23]. The possibility of inducing pathological conditions by using TGF-β2 and dexamethasone was shown, and the adverse effect of benzalkonium chloride, an eyedrop preservative, was revealed by induced inflammatory chemokines and inhibited MMP activity [23]. None of these studies took into account the gradient structure of the HTM to recapitulate in vivo tissue morphology. Biomimetic HTM scaffolds that could better reproduce native morphological features and help to better understand structure-function relationship are still needed [24].

Multiphasic or gradient scaffolds for in vitro engineering of tissues can be fabricated by electrospinning techniques [25]. In this method, layers of fibrils with controlled dimensions and at controlled density can be deposited [26, 27]. Using melt electrowriting (MEW), a marriage between electrospinning and 3D printing, gradient scaffolds with aligned fibers can be obtained. In MEW, fibers with thickness in the micrometer range are deposited from the melt with the aid of electrical voltage using a robotic stage [28, 29]. Well-defined, highly porous architectures can be fabricated with flexibility in the dimensions and spacing of the fibers and the number of laydown layers [30]. MEW has been applied to print thermoplastic polymers like poly(caprolactone) (PCL) into structures with different pore architectures, including squared [30-34], rectangular [33], rhombus [34], dodecagon [30], triangle [30] or sinusoidal [35, 36]. When used as scaffolds for cell culture, cell growth has been demonstrated to depend on the fiber size [32] and pore geometry [37]. Gradient and multiphasic PCL scaffolds with pore sizes in the range of 250 to 750 µm and 10 µm thick fibers have been fabricated and applied for bone regeneration [38]. PCL scaffolds with pore sizes in the range of 125 to 250 µm and fiber diameters from 4 to 25 µm were used to culture human adipose-derived stem cell spheroids [31]. Scaffolds with pore size 100 to 400 µm and fiber diameter of ca. 10 µm were used for cartilage regeneration [39, 40]. MEW has been used for skin [41], cardiac [33, 36], and ligament tissue engineering [35], and biomimetic designs of tympanic membrane [42] and cartilage [39, 40].

In this study, MEW is applied to obtain gradient scaffolds of poly(caprolactone) that mimic the morphological characteristics of native HTM. This is the first report on MEW application in ocular tissue engineering. The methodology to prepare scaffolds containing up to 88 stacked layers of fibrils with a graded pore architecture is described. The topology, porosity, and mechanical properties of the scaffolds as a function of the design are characterized. The morphology of the primary HTM cells cultured on the scaffolds is described as a function of the scaffold’s geometry. The results indicate mechanical properties of the scaffolds matched those of natural HTM, with the cells maintaining the phenotype of native HTM cells and infiltrating the scaffolds. The utility of MEW to mimic complex morphological features of small-scale gradient natural tissues is demonstrated. This study paves the way to develop functional biomimetic high adequacy in vitro models that will allow to develop a detailed understanding of the structure-function relationship in native HTM.

## 2. Results and Discussion

### 2.1. Scaffold Design and Printing

Inspired by the distinctive structure and pore size of the consecutive layers of the native HTM, three different scaffolds were designed (PCL 16, PCL 60, and PCL 12; the numbers indicate the number of deposited layers). The targeted dimensions for printing were a fiber diameter of 10 μm, recapitulating typical trabecular beam size, and 80 % porosity, with effective pore sizes decreasing in the order PCL12 < PCL 60 ≤ PCL 16. Scaffolds PCL 16 and PCL 60 **(Figure 1)**, containing 16 and 60 printed layers, respectively, were constructed by a periodically repeated square mesh pattern (with 200 µm interfiber spacing). In PCL 16, the consecutive square meshes were shifted by ½ of the period (100 µm) in x and y axis; in PCL 60 the consecutive square meshes were rotated by 30°, and after every 3 square meshes a shift of ½ of the period (100 µm) in x and y axis was applied. In PCL 12 (**Figure 1**), with 12 printed layers, fibers in each layer were printed with smaller interfiber distance (100 µm) and were rotated by 15° to achieve smaller pore sizes than in previous designs. A PCL full scaffold was also fabricated (**Figure 1**) by superposing the three previous designs to reconstruct the multilayer HTM structure.

**Figure 1.**
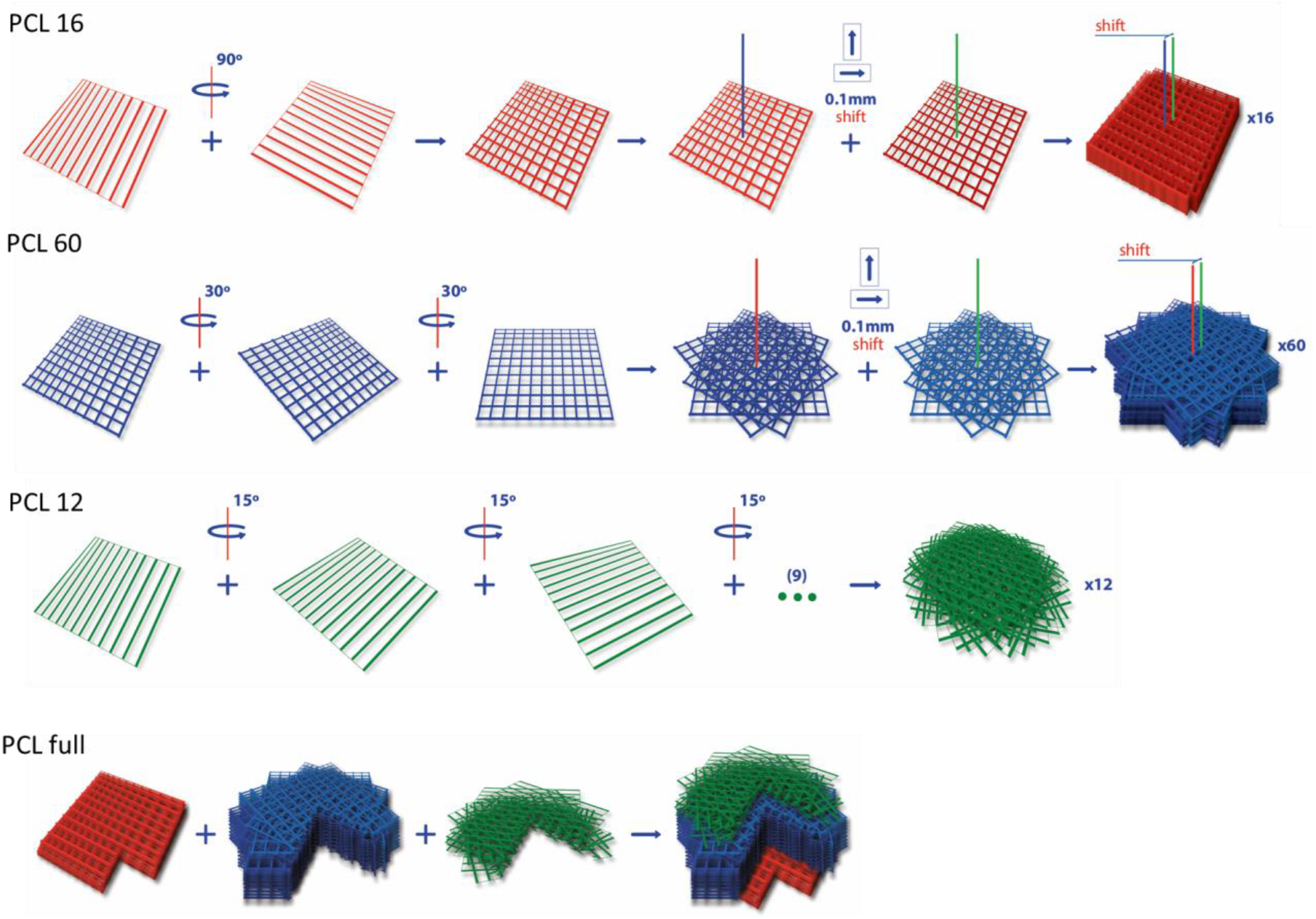
Scaffold design and rotations/displacements performed in the printing steps. PCL full structure shown with cross-section.

Medical grade PCL ca. 100 000 kDa (see **Table S1**) was selected as electrowriting ink due to its good processability [43, 44] and biocompatibility [45]. Printing parameters were optimized to achieve the best shape fidelity, bedside PCL 12 design, as discussed later. Scaffolds contained superposed layers of 25 × 25 mm^2^ area **(Figure 2A)** and were printed layer-by-layer. The layers contained aligned fibers with 10-12 µm diameter **(Table 1)** and spacing of 200 µm (PCL 16 and 60) or 100 µm (PCL 12). 200 µm was the minimum interfiber distance that we could achieve with high fidelity, in agreement with previous work on MEW with PCL by other authors [35]. Shorter interfiber distances lead to lower precision in the fiber deposition due to the charge built up at the collector plate and the consequent alterations in the electrical field [46-48]. PCL 16 and PCL 60 scaffolds displayed straight and parallel fibers, albeit with a few distortions in the printed fibers (indicated by white arrows in **Figure 2B**). In PCL 12, in order to reduce pore size, a smaller interlayer rotation (15°) was used, and a 100 µm interfiber distance was attempted, though this was at the cost of a more irregular material flow and bending of the deposited fibers. Additionally, printing below critical translational speed was used intentionally to introduce fiber crimps and further decrease pore sizes.

**Table 1.** Features of scaffolds printed with different designs: height, fiber diameter, theoretical and measure porosity, and water contact angle.

| | Scaffold height ($\mu\text{m}$) | Fiber diameter ($\mu\text{m}$) | Porosity (%) | | Contact Angle ($^{\circ}$) | Compression modulus E (kPa) | | Tensile modulus E (MPa) | |
| --- | --- | --- | --- | --- | --- | --- | --- | --- | --- |
|  |  |  | theoretical | measured |  | dry | wet | dry | wet |
| **PCL 12** | 125±10 | 10.0±0.6 | 92.15 | 86.8±1.9 | 130±2 | 11.2±3.3 | 5.6±1.9 | 13.0±1.7 | 11.1±1.1 |
| **PCL 16** | 140±19 | 11.8±1.4 | 96.07 | 91.2±1.6 <sup>a</sup> | 130±4 | 63.8±79.9 | 66.4±87.6 | 7.2±2.1 | 5.6±1.8 |
| **PCL 60** | 299±20 | 10.2±0.8 | 96.07 | 84.7±0.4 | 126±1 | 87.9±75.6 | 216.4±175.7 | 7.6±0.9 | 7.3±1.0 |
| **PCL full** | 506±18 | 11.9±1.3 | 95.54 | 84.2±1.5 | 126±8 | 358±235 | 35±17 <sup>b</sup> | 6.9±1.1 | 6.7±1.4 |

<sup>a</sup> statistically different from all other scaffolds
<sup>b</sup> statistically different from dry scaffold

**Figure 2.**
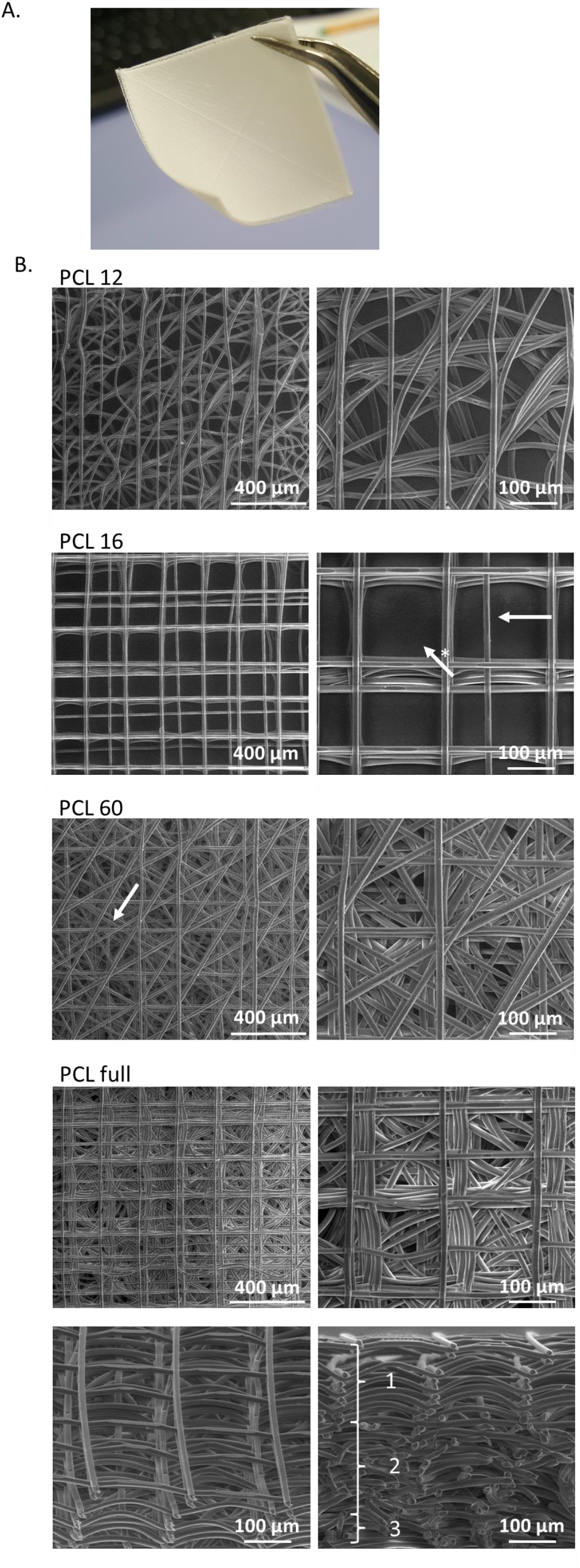
Images of printed scaffolds. Macroscopic view of printed PCL full scaffold (A). SEM top view images at 200x (left) and 500x (right) magnification. A few structural defects are highlighted: white arrows show example of not precisely positioned fibers, white arrow with (*) shows a missing vertical fiber according to the design. The bottom row presents the tilted and cross-section view of PCL full at magnification 500x. Three distinctive layers are marked in white (B).

The thickness of the printed scaffolds (**Table 1**) varied between ca. 125 µm (PCL 12) and ca. 500 um (PCL full). The total thickness of scaffolds was smaller than the sum of the consecutive printed layers due to the layers merging, assuring proper connectivity and minimizing delamination. The measured porosity, based on the previously reported apparent density approach [49], was 84 % - 91 %, with the order of PCL full ≈ PCL60 < PCL 12 < PCL 16 (see **Table 1**), whereas the theoretical porosity, for ideal scaffolds with 10 μm diameter strands and no merging between consecutive printed layers, was calculated as 92 % – 96 %. The measured porosity was smaller than the theoretical one for all the scaffolds due to the merging between layers, resulting in material densification, and fiber diameter exceeding 10 μm in the case of PCL 16 and PCL full scaffolds (**Table 1**). The smallest difference between theoretical and measured porosity was detected for PCL 16 scaffolds, indicating the highest printing fidelity for this design. On the contrary, the highest deviation between theoretical and measured values was observed for PCL 60, which we assigned to the most pronounced merging of the consecutive printed layers due to the highest number of them, resulting in the highest scaffold weight. Note that despite the porosity % being in the same range for PCL 12 and PCL 60, the pore sizes for PCL 12, meaning the distance between adjacent fibers, were smaller due to the designed 100 µm interfiber distance.

### 2.2. Wetting Properties and Stability of the Printed Scaffolds in Watery Media

The wetting properties of the scaffold were studied by measuring the water contact angle. Values between 125° and 130° were observed (**Table 1**), indicating hydrophobicity of the scaffold’s surface as a consequence of their surface structure. Note that the PCL film has a contact angle below 90 ° [50]. Importantly, the scaffolds immersed in an aqueous solution were wetted instantaneously in the case of PCL 12 and PCL 16, and slower in the case of many-layered structures (PCL 60, PCL full). The facilitated wetting of PCL 12 and PCL 16 can be explained by easier removal of the air pockets in thinner scaffolds during immersion [51]. All scaffolds remained stable during immersion in a cell culture medium for 14 days. No delamination or disintegration was observed. This result indicates good interfibrillar adhesion between the printed fiber and the one below. Note that the degradation time of PCL in water is 12 months [52].

### 2.3. Mechanical Properties

Uniaxial compression tests of the scaffolds in the static mode were performed. The strain-stress curves (**Figure 3A-B** and **Figure S2**) show two regions of different slopes: an initial region of lower slope at strains <40% (PCL 60 and PCL full) and <60% (PCL 12 and PCL 16), which was followed by a densification region with higher slope [53]. Based on the strain–stress curves, the compression behavior of PCL full seems to be dominated by the compression behavior of the PCL 60 region, which is the main constituent regarding scaffold volume (60 layers of 88 layers in total). The elastic modulus of the printed scaffolds was extracted from the initial slope of the curve (<10% strain). The obtained moduli are in the range of 6 – 360 kPa (see **Table 1**), with the values showing the trend: PCL12 < PCL 16 < PCL 60 <PCL full for the dry samples (no statistical significance). This subtle effect correlates to the scaffold porosity (PCL full ≈ PCL 60 <PCL 16 < PCL12) and the number of printed layers (PCL12 < PCL 16 < PCL 60 <PCL full), and could be explained by more extensive merging between the layers leading to the increased connection at the nodes. The compressive modulus values show a similar trend to previous studies on highly porous PCL scaffolds, suggesting that the modulus is related to porosity and less to fiber alignment or design geometry [53]. The compression measurements were performed in a dry and wet state (note that PCL shows limited swelling when immersed in water [54]). There were no statistical differences between wet and dry samples, except for PCL full, which revealed decreased modulus value after 24 – 48 hours of incubation in PBS at RT. This could be due to differences in the water uptake of the different constituting layers, resulting in local delamination or defects in connection points [54]. The compression moduli values are in agreement with previous reports. For multilayered PCL scaffolds printed with MEW with a square-based design, ca. 20 μm fiber diameter, 200 μm interfiber spacing and 1 mm height, compression moduli of ca. 14 kPa [55] or slightly below 1 MPa [39] were reported. The ranges of compression moduli obtained in the study are close to those of collagen-based scaffolds used for culturing HTM cells by other authors (7kPa [20]). The compression modulus of HTM in healthy and glaucoma HTM has been reported to be 4 and 80 kPa, respectively, yet measured by AFM [12]. Other studies reported only 1.4-fold increase in the storage modulus of glaucomatous HTM [14, 15]. The scaffolds withstood strains up to at least 60% before failure (**Figure S3**). For PCL 12 and PCL 16 the failure was not recorded due to the measurement limitation for thin samples. The failure behavior in PCL full was dominated by the thickest, PCL 60 layer.

**Figure 3.**
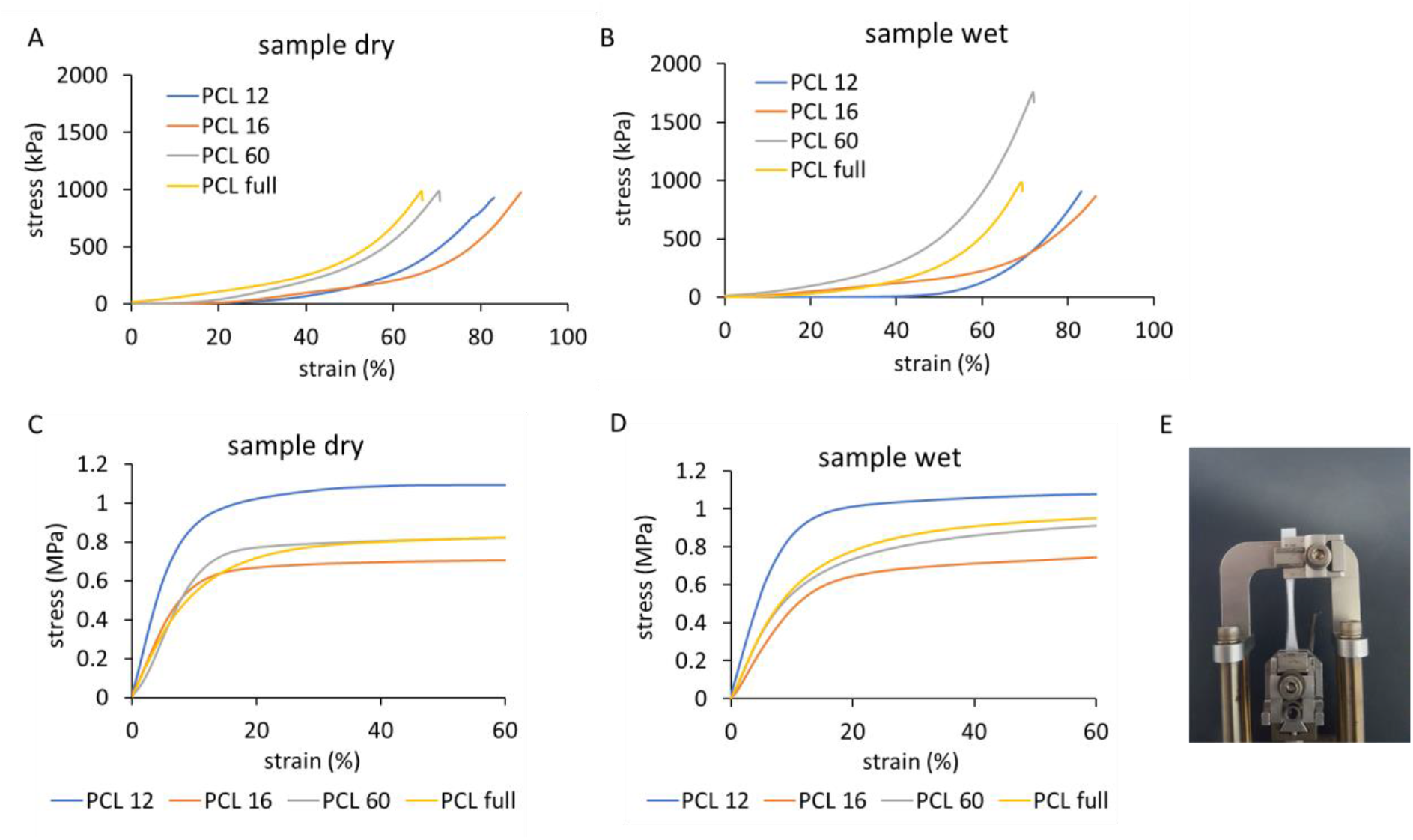
Mechanical testing of different scaffolds. Stress-strain curves of the scaffolds obtained in a static compression test of dry (A) and wet (B) samples; and in a tensile test of dry (C) and wet (D) samples. PCL 60 sample in the end of the test, after ca. 300% elongation (E).

The stress-strain curves in the uniaxial tensile test of the scaffolds are shown in **Figure 3C-D** and **Figure S4-S5**. An initial linear region with a steep slope was followed by a long plateau region. No failure was observed below 300% elongation, which was the extension limit of our equipment (**Figure 3E**). The tensile modulus was in the range of 5.6 - 13 MPa for all the designs (see **Table 1**). These values are in agreement with previously observed for square-based scaffolds [33, 39], where the range of a few MPa was reached. The modulus of PCL 16, PCL 60, and PCL full was similar; the slightly higher modulus of PCL 12 might be associated with the lower porosity (highest material volume fraction). In PCL 16, fibers elongated in the stretching direction without visible fiber break, whereas in PCL 12, PCL 60, and PCL full failure of consecutive, single fibers was observed during the experiment (**Figure S5**). No variation in the tensile modulus was observed with the humidity of the sample.

Different values for Young’s Modulus of dissected HTM have been reported. Human HTM segments measured by uniaxial tensile test showed a Young’s Modulus of 515 ± 136 kPa, while porcine HTM showed 25 kPa [56]. In a different report, 12.5 MPa were measured for glaucomatous human HTM and 42.6 MPa for normal tissue [57]. The results obtained here seem to be within the range relevant for physiological studies.

### 2.4. Cell culture studies

The ability of the printed scaffolds to support the culture of primary HTM cells was tested in viability and metabolic activity tests. HTM cells were seeded on the scaffolds. We performed a live/dead assay to assess cell viability, followed by fluorescent imaging. In PCL 12 and PCL 16 samples, the stained cells were visible through the scaffolds, whereas for the PCL 60 and PCL full scaffolds that contain a high number of layers, only cells in the more superficial layers were available for examination and included in the analysis. Cell viability varied between 60% and 90 % at day 1, day 8, and day 14 after seeding (**Figure 4A**). PCL 60 and PCL full show a similar trend, which can be explained by the fact that PCL 60 constitutes the main part of PCL full and that the cells after seeding could penetrate the PCL 60 layer. Relatively lower viability was detected in those scaffolds compared to PCL12 and PCL 16. The drop in viability for all the samples on day 14 is attributed to the high cell density achieved at that time point. The large error in the measurements is due to imaging difficulties as a consequence of the reflection of the light by the scaffold.

**Figure 4.**
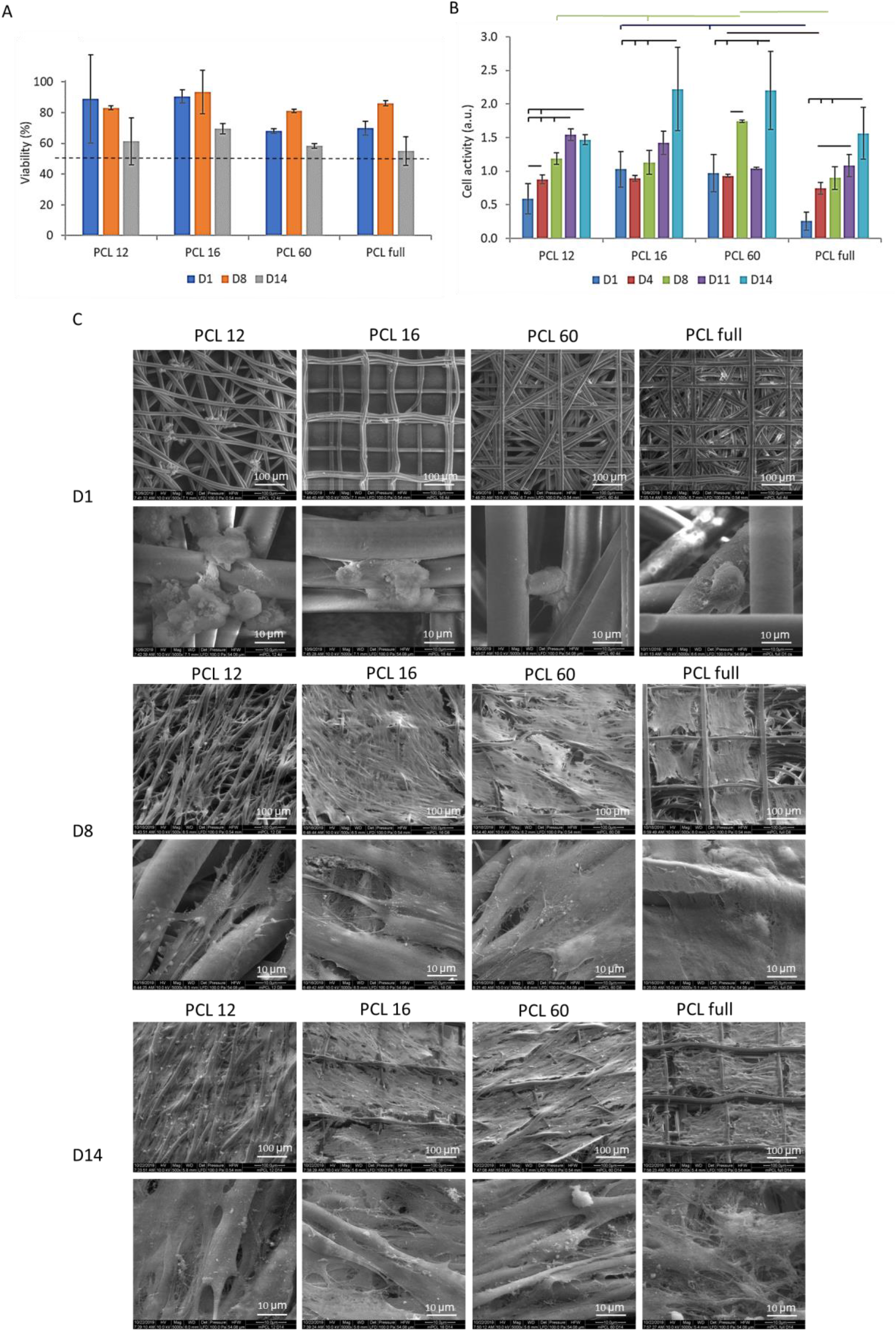
Biological evaluation of the cells growing on different scaffolds. A. Viability of the cells based on live/dead assay staining. Threshold of 50% viability marked by the broken line. B. Cell proliferation based on AlamarBlue activity assay. C. SEM images of scaffolds with cells at day 1, day 8 and day 14 at 2 magnifications: 500x (top; scale bars: 100 µm) and 5000x (bottom; scale bars: 10 µm).

Cell proliferation within the substrates was estimated by the AlamarBlue assay (**Figure 4B**), which tests the metabolic activity of the cells. Metabolic activity was observed across all scaffolds, and it increased with culture time, indicating cell proliferation regardless of the design.

#### 2.4.1. Cell Morphology and Distribution on the Scaffolds

The morphology of HTM cells on the scaffolds on days 1, 8, and 14 after seeding was studied by scanning electron microscopy. On day 1 (**Figure 4C**), cells accumulated at the nodes or fiber crossings of the scaffolds and had a rounded morphology. On day 8, cells spread out on the scaffolds, revealing a typical spindle-like shape (native HTM cell morphology) and overlapping processes [20]. However, different morphologies and coverage areas were observed depending on the scaffold design. Cells on PCL 16, PCL 60, and PCL full (which was seeded on the PCL 16 surface) spread out in all directions and spanned the fibers of the scaffold’s surface, filling the interfiber space, and formed a dense cell layer. The covered area by the cells was larger in PCL 16 and 60 than in PCL full. In contrast, cells on PCL 12 scaffolds elongated along the fibers and, in some cases, bridged adjacent fibers but did not form continuous cell layers of appreciable area. On day 14, PCL 12 and PCL 60 scaffolds were covered by a uniform and dense cell layer, whereas the cell layer on PCL 16 and PCL full had some empty areas (see **Figure S6**). The SEM investigation (see **Figures S7** and **S8**) and the nuclei distribution tracked with confocal microscopy **(Figure S9)** confirmed that cells could infiltrate all the scaffolds. We attribute the differences in the cell density inside the scaffolds observed with both imaging techniques to the light reflection on the PCL mesh that hinders fluorescent imaging of the cells in the multilayer scaffolds. The most uniform infiltration was detected for thinner scaffolds (PCL 12 and PCL 16). The smaller pores sizes facilitated bridging of the fibers at earlier time points, and confluent layer formation on the surface of the scaffold. For planar SU-8 scaffolds reported before, it was observed that the cells have difficulty growing on pore sizes bigger than 15 μm [19]. For those scaffolds [19] and 3D hydrogel-based scaffolds produced by freeze-drying [20], limited cell penetration into the pores was observed, whereas in our models, the cell infiltration was significant. In the follow-up study, the authors have shown that the cells seeded on freeze-dried hydrogel-based scaffolds with large pore sizes (in the range of 200 μm) proliferate more than on the samples with small pores (in the range of 20 μm), most probably thanks to the higher surface area available for cells growth and easier nutrients and oxygen transport. The non-aligned pores were also more beneficial for cell growth than the aligned ones due to the alternative routes for cell proliferation and migration [21].

#### 2.4.2. Cellular identity/maintenance of the phenotype

HTM cells in the natural tissue have elongated cell shapes bridging multiple adjacent fibers, elongated nuclei, and reveal specific markers [8]. To further analyze the phenotype of HTM cells within the scaffolds, cells were stained with DAPI, Phalloidin, and anti-αβ-crystallin to reveal nucleus elongation, the disposition of actin fibers, and expression of HTM cell-specific marker, respectively.

The nucleus shape in cells on the different designs based on DAPI staining was estimated after 14 days of culture. The Aspect Ratio (AR) parameter was calculated as a ratio of nucleus length to width (an example of image preparation for quantifications is provided in SI, **Figure S10**). The nuclei of the cells were elongated (Aspect Ratio, AR ≥ 1.5: 1.7 ± 0.4 for PCL 12, 1.7 ± 0.5 for PCL 16, and 1.5 ± 0.4 for PCL 60, see also **Figure 5**) with greater elongation visible for PCL 12 and PCL 16. The native HTM cells in the in vivo conditions typically reveal elongated nuclei [8]. The AR reported in previous studies was 1.1 for 2D culture on a porous membrane (insert) and 2.0 for optimized SU-8 photoresist membrane with pores of 12μm.

**Figure 5.**
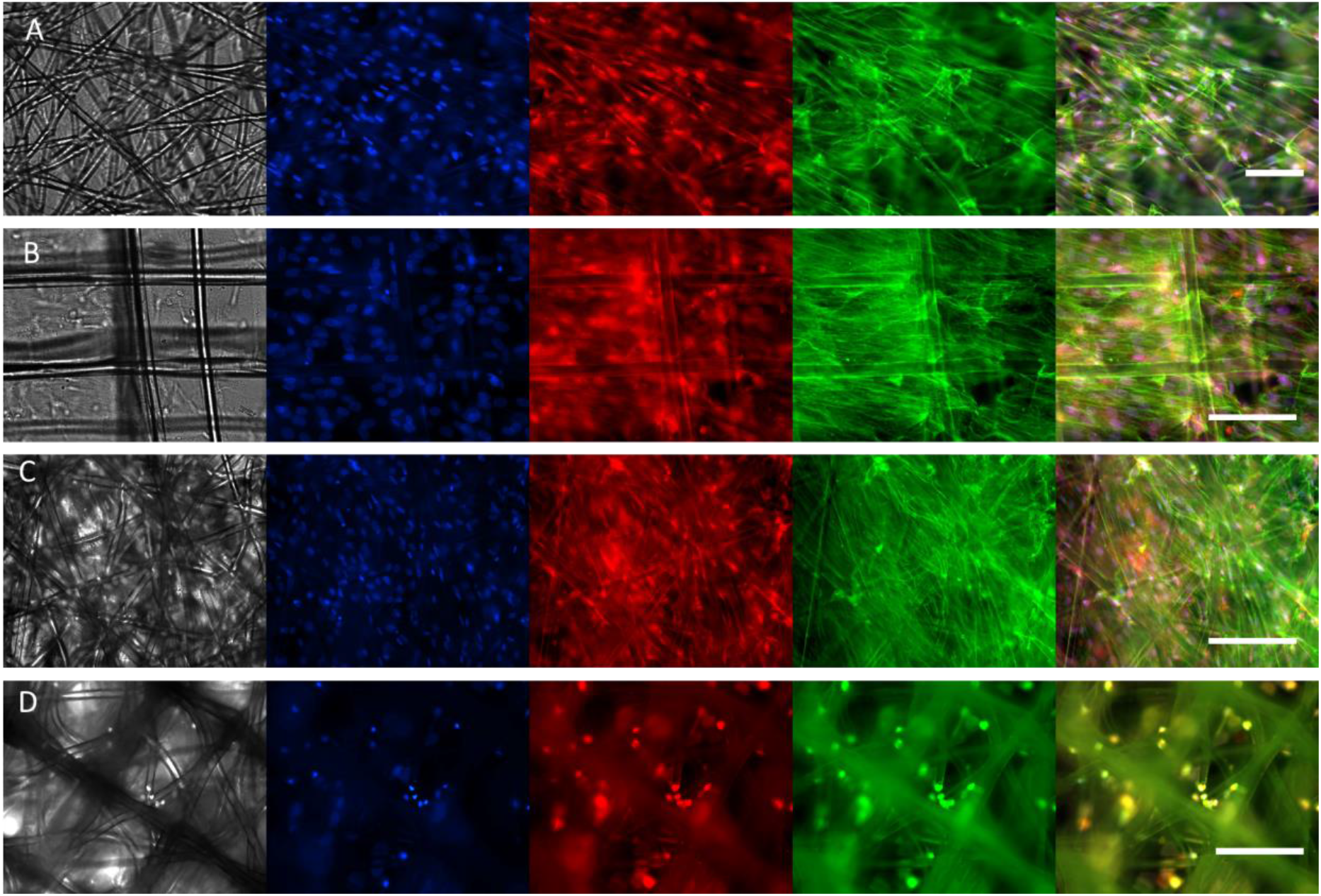
Fluorescent images of the cells cultured for 14 days on different scaffold types: PCL 12 (A), PCL 16 (B), PCL 60 (C), PCL full (D). The first channel represents bright-field (in gray), the second channel DAPI nuclei staining (blue), the third channel ab-crystallin (red), the fourth channel phalloidin actin staining (green), and the last tile is a merged representation. Scale bars 100 µm. Note that due to the light reflection, the light imaging of the cells in multilayer scaffolds is impeded.

Actin staining after 14 days of culture showed elongated structures in the cells cultured on all the scaffolds, besides PCL full (**Figure 5** and **S11**), indicating that cells are attached to the scaffold and are able to build cytoskeletal actin fibers. Due to the reflection of the light by the thick, nontransparent scaffold, the cellular cytoskeleton and spreading of the cells on PCL full was difficult to observe based on actin staining and the clear alignment observed in other studies [8] was difficult to prove. Nevertheless, SEM images indicate the presence of stretched cells, bridging adjacent fibers in all printed designs. αβ-crystallin was expressed by HTM cells on all the scaffolds. αβ-crystallin is a characteristic marker of the JCT region in HTM [7] and is not expressed in conventional 2D cultures [19]. Therefore, we concluded that the MEW-printed 3D HTM model revealed improved properties, leading to the appearance of the key in vivo-like HTM characteristics.

The obtained data show that the printed PCL scaffolds were suitable for the culture of HTM cells and maintained the native phenotype of JCT cells, including the spreading and formation of cell layers, with cell-cell interfaces between adjacent cells. The native JCT is covered by 2-5 discontinuous cell layers, with the cells making a satellite connection with the endothelial cells lining the Schlemm Cannel [5]. The cells in native UM and CM regions have more rounded shapes, whereas cells in JCT region adopt elongated shapes [3, 24]. In our scaffolds, the JCT phenotype seems to be reconstituted. Changes in the printed designs, such as more open porosity (less dense structure) and a smaller number of the printed layers in the middle zone, might allow reconstruction of the phenotype of HTM cells in the UM or CM layers.

## 3. Conclusion

This study demonstrates the feasibility of in vitro reconstruction of some features of the human trabecular meshwork using MEW scaffolds of PCL, i.e. porosity gradient and native trabecular beam size. It is the first report on MEW application in ocular tissue engineering. Scaffolds with three different designs were printed in an effort to build the three distinctive matrix layers of native HTM, and one combined design containing a stacking of the three layers with a gradient of morphologies and porosities. Printed scaffolds had relevant mechanical properties, were stable (no delamination effect) during 14 days of cell culture, and supported 14 days of culture of primary HTM cells. Scaffold design influenced cell morphology: the thinner scaffolds showed better cell infiltration, and the smaller pores sizes facilitated cell elongation along the fibers; the bridging of multiple adjacent fibers at an earlier time point (8 days), and confluent and more uniform layer formation on the surface of the scaffold at the later time point (after 14 days). HTM cells on the scaffolds showed elongated nuclei, a well-developed actin cytoskeleton, and revealed a specific marker observed only in 3D culturing conditions, which are features characteristic of HTM cells in the natural JCL layer of the native human trabecular meshwork tissue. This study opens the way to produce biomimetic functional HTM engineered scaffolds, to improve understanding of structure-function relations in the small-scale gradient tissues. Permeability studies would be necessary to validate the printed models in further work.

## 4. Methods

### 4.1. Fabrication of MEW Scaffolds: Design and Printing parameters

Melt electrowriting was performed using a 3D Discovery printer (RegenHu, Switzerland) integrated into a safety cabinet (sterile conditions). The scaffold designs were programmed using BioCAD™ software (RegenHu, Switzerland). Four scaffold types with a different number of deposited layers (12, 16, 60, or 88), resulting thickness (125 to 500 μm), and fiber orientation between consequent layers (15 °, 30 °, or 90°) were prepared. They were named PCL 16, PCL 60, PCL 12, or PCL full, where the numbers indicate the number of deposited layers. The term “full” refers to the gradient scaffold in which the three designs (PCL 16, PCL 60, PCL 12) were printed consecutively (for visual presentation, see Figure. 1). 25 × 25 mm^2^ layers were printed containing parallel strands of ca. 10-12 μm diameter. The interline spacing was set to 0.2 mm for PCL 16 and PCL 60, and to 0.1 mm for PCL 12. In PCL 16, two consecutive squares rotated by 90° were printed first, followed by an 8-fold repetition of this pattern with a 0.1 mm shift in the x and y axis (16 layers in total). For PCL 12, twelve consecutive squares were printed, maintaining a rotation of 15° for each layer. For PCL 60, three consecutive squares were printed with 60° rotation each, and this pattern was repeated 20 times with a 0.1 mm shift in the x and y axis (60 layers in total). For PCL full, the three described patterns were printed continuously in the order PCL 16, PCL 60, and PCL 12, to mimic the gradient structure of the native HTM. Medical-grade PCL PURASORB PC 12 with Mn = 80,000 g/mol (Corbion Inc, Netherlands) was used for printing. The following printing parameters were applied: 90°C cartridge temperature, 60 kPa pressure, 10 mm/s printing speed, 10 kV voltage, 3 mm nozzle distance from the collector, 22G (0.4 mm) nozzle size. The material was heated to 90°C for two hours before the printing started and refreshed in the cartridge after every three heating cycles. Before printing, the jet was stabilized by printing a sample scaffold with the final printing parameters for three minutes. The material deposition was performed onto the collector covered with an A4 inkjet transparency film to facilitate scaffold removal. The printing process was monitored with a high-speed camera integrated with the printing head.

### 4.2. Imaging and Morphological Characterization of the Scaffolds

The printed scaffolds were imaged with a stereomicroscope SMZ800N (Nikon, Germany) using bottom illumination. The width of the printed strands was measured with the integrated NISElements D (Nikon, German) software at a minimum of nine different locations in the scaffold. Four independent scaffolds of each type were printed and analyzed.

The scaffolds were also imaged by scanning electron microscopy (SEM – FEI Quanta 400 FEG). For this purpose, a sample of 5 × 5 mm^2^ was cut with a blade and fixed to an aluminum holder using double-sided adhesive carbon tape. For the analysis, top view and cross section images were taken. Secondary Electron (SE) imaging was performed at 10 kV accelerating voltage in low vacuum mode (p = 100 Pa water vapor, Large Field Detector – LFD, dwell time 1.5 µs, spot size 3).

To measure scaffolds porosity, the scaffold dimensions and weight were measured. Scaffold height was measured using a rheometer (gap value at axial force increased to ca. 0.05N). Porosity was calculated according to the apparent density approach [49], by the equation: p = (1-m/V)/ρ_PCL_ * 100%, where p = porosity, m = scaffold mass, V = scaffold volume, and ρ_PCL_ = density of PCL (1.145 g cm^-3^). Measurements were performed on four samples per scaffold type. The measured porosity was between 80 % and 90 %. The expected porosity values were calculated for each scaffold type based on the volume occupied by printed strands (approximated as ideal cylinders) with 10 μm diameter and with the assumption that the consecutive layers are not merging but touching at one point. Consequently, based on the design the porosity for PCL 16 and PCL 60 was 96.07%, 96.07%, 92.15% and 95.54 % for PCL 16, PCL 60, PCL 12 and PCL full, respectively.

### 4.3. PCL Characterization

The molecular weight of PCL as purchased, after melting and heating for two hours and after printing, was measured by gel permeation chromatography (PSS GPC-MALLS, Germany) using the refractive index for detection. The samples were dissolved in Tetrahydrofuran (THF) at 30°C for 20 minutes at a concentration 2 g/l and filtrated with a 0.45 μm PTFE membrane syringe filter. 20 μl of the filtered sample were used for HPLC-GPC analysis using the following conditions: 1 ml/min flow rate, 40°C, 66 bar. For calibration 2 g /l solutions of Polystyrene standards (PSS-Polymer, Germany) were used.

FT-IR analysis of PCL as purchased, after melting and heating for two hours and after printing, was performed (Bruker TENSOR 27 equipped with Specac’s ATR Golden Gate, USA). Measurements were taken in triplicates; the most representative spectra are shown.

The thermal properties of the PCL (melting temperature (*T*_*m*_) and crystallinity) samples were measured by differential scanning calorimetry (DSC1 STAR^e^ System, Mettler Toledo). Two heating/cooling cycles at 5°C/min were performed; reported results correspond to the second heating-cooling cycle. The crystallinity fraction was calculated from the melting enthalpy taking 139.5 J/g as the corresponding enthalpy to 100% crystallinity [58].

GPC, FTIR, and DSC analyses of samples as received, after melting, and after printing were performed to identify possible changes in the polymer during processing. No significant changes were observed (Table S1 and Figure S1).

### 4.4. Characterization of Wetting Properties of the Scaffolds

Contact angle measurements on the scaffolds were performed using an OCA20 (Dataphysics instruments GmbH, Germany). Measurements were performed on three different scaffold samples at room temperature, using 1 μl water droplet. In PCL full, the droplet was seeded on the side corresponding to the PCL 16 design. The images were captured using a high-speed camera. Movies of sample wetting at room temperature were captured by dipping the scaffold into poly-L lysine solution while imaging with a stereomicroscope (Olympus SZX16) equipped with an Olympus SC50 CCD camera and Software Olympus Stream Basic 1.9.4.

### 4.5. Characterization of Mechanical Properties of the Scaffolds

Static compression and uniaxial tensile tests were performed in the wet and dry states. Wet samples were incubated for 24 – 48 hours in PBS solution. Measurements were performed on three samples. On the final plots representing compressive modulus data, the most representative curves are depicted. All the obtained curves are presented in SI.

For the compression test in static mode, a TA Rheometer (DHR3, TA Instruments, USA) with parallel plate geometry was used. Round samples of 6- or 8-mm radius were cut with a sharp plunger at different spots of printed scaffolds. Prior to experiment initialization, the samples were placed at the center of the bottom plate using tweezers. For wet samples, PBS solution was added to the bottom plate around the scaffold. All experiments were performed at room temperature. Prior to measurement initialization, the upper plate of the Rheometer was manually driven to get into contact with the scaffolds (ca. 0.2 N axial force), and the sample was compressed at a speed 0.2 µm/s. The initial compressive modulus was calculated from the slope of the linear part of the stress vs. strain plot, in the range of 1 - 10% strain, using TIROS software (DHR3, TA Instruments, USA). Measurements performed below 1% strain showed high variability. One sample (out of three) measured for PCL 12 revealed the fit of linear regression R < 0.8 and was discharged from the calculation of the modulus value. For all other samples, linear regression fit gave the R > 0.8. The maximal stress and strain values were determined from the peak of the stress-strain curve.

Uniaxial tensile tests were performed using a Q800 DMA (TA Instruments, USA). The printed scaffolds were cut as 5 mm broad stripes (25 mm long), with the longer axis parallel to fibers deposited in the first printed layer. The load was applied along the longer axis. Measurements were performed in the force range of 0.01 to 1 N at a constant force ramp 0.1 N/min, at room temperature, and with preload force of 0.01 N. The distance between clamps was 5-6 mm. A 30 s temperature equilibration time was applied prior to the initialization of the experiment. Three samples were tested for each scaffold type to a maximum extension of 15 mm (limit of the machine). The effective elastic modulus was calculated from the slope of the linear region of the stress vs. strain curve. Ultimate stress and ductility were calculated from the plots, based on the maximum stress and strain recorded, and presented in the Supplementary Information (**Figure S5**).

### 4.6. Cell Experiments

Primary Human Trabecular Meshwork cells isolated from the juxtacanalicular and corneoscleral regions (P10879, Innoprot, Spain) were cultured according to the provider’s protocol. In short, the culture was setup in a 0.1 % Poly-L-lysine coated T75-flasks at a seeding density of 5,000 cells/cm^2^, in RPMI 1640 medium (Gibco, 61870-010) supplemented with 10% FBS (Gibco, 10270), 1 ng/ml fibroblast growth factors, 200 mM L-glutamine and 1% penicillin/streptomycin (Invitrogen), at 37°C in a humidified atmosphere of 5% CO_2_. The medium was refreshed every 2-3 days. When 90% confluency was reached, the cells were subcultured.

5 mm wide stripes cut from 25×25 mm^2^ scaffolds and spherical samples (6 mm diameter) cut from 6×25 mm^2^ scaffolds were used for the experiments. All scaffolds were sterilized prior to cell seeding by two immersions in 70% ethanol for 20 minutes, followed by washing twice with PBS and soaking over weekend in 70% ethanol. Afterward, samples were incubated in sterile PBS solution for 1.5 h at 37°C and finally washed twice with sterile PBS.

Cells were then seeded on the 4 scaffold types (for scaffold PCL full, cells were seeded on PCL 16 side). Prior to seeding, scaffolds were placed in cell culture plates: spherical scaffolds in 96-well plates and the stripes in 6-well plates and fixed to the bottom of the plate by a home-made plastic ring (prepared from Thincerts™ (Greiner Bio-One, Germany) by removing the bottom membrane). Scaffolds were not coated with adhesive proteins. HTM cells at passage 5 were suspended in a culture medium at 1 million/ml and seeded on top of the scaffolds (40 μl onto small scaffolds, 50 μl onto stripes), giving the final cell density of 40 000/cm^2^. After 1-2 hours of incubation, an additional medium was added (200 μl per well to 96-well plate, and 3 ml per well to 6-well plate). The seeded scaffolds were kept in culture for 14 days, with the cell culture medium changed every 2-3 days. Small scaffolds were used for the cell metabolic activity test, performed with an AlamarBlue assay. 5mm-width stripes at day 1, day 8, and day 14 were cut with a sterile blade to be used for viability assay, immunostaining and SEM investigation or further culture.

#### 4.6.1. Cell Viability

A live/dead cell viability assay was performed following the manufacturer’s protocol (Sigma-Aldrich). Briefly, PBS solutions of 20 μg /ml propidium iodide (PI), to stain dead cells in red (excitation/emission ≈535/617 nm), and 6 μg / ml fluorescein diacetate (FDA), to stain live cells in green (excitation/emission ≈490/515 nm), were prepared. Samples at day 1, day 8 and day 14 were removed from the medium, placed in a fresh well plate, and incubated with 100 μl – 250 μl of staining solution for 10 min. After 2x washing with PBS, cells on the scaffolds were imaged with a PolScope fluorescent microscope (Zeiss, Germany). Images were analyzed with ImageJ software. After brightness and contrast adjustment, cell counting was performed with the function ‘find maxima’. The cell viability was calculated using the following equation: % Viability = (n° of live cells / total n° of cells) x 100.

#### 4.6.2. Cell Proliferation, Nuclei Shape and Scaffolds Infiltration

Cell proliferation was quantified using the Alamar Blue assay (Invitrogen). Samples on days 1, 4, 8, 11 and 14 were transferred to a fresh well plate, and the Alamar Blue reagent was added in a 1:10 ratio to the culturing medium. After 3.5 h, the scaffolds were removed, and the absorbance of the medium at 570 nm was analyzed with a Multi-Detection Microplate Reader Synergy HT (BioTek Instruments; Vermont, USA). For each scaffold type, three independent samples were analyzed. Results were normalized to the control (cells seeded on the bottom of 96-well plate at day 0 at the density of 40000/cm^2^).

Samples on day 14 were fixed with PFA 4% w/v for 10 min, followed by three washes with PBS. After permeabilization with 0.5% w/v Triton-X 100 (TX) for 10 min, and PBS wash, cell nuclei were stained with 1:1000 DAPI (4′,6-Diamidino-2-Phenylindole, Dihydrochloride, ThermoFisher) in PBS, and washed with PBS again. Imaging was performed with Nikon Ti-Eclipse (Nikon Instruments Europe B.V., Germany) with a Sola SE 365 II (Lumencor Inc., Beaverton, USA) solid-state illumination device and an Andor Clara CCD camera. Three independent samples imaged at 20x magnification were used to analyze nuclei parameters: length and width, and the ratio of length to width, farther denoted as AR (Aspect Ratio). The pictures were processed with ImageJ Software by color threshold adjustment, followed by watershed processing and the use of the “analyze particles” function (size limit 50 – 150 µm). Scaffolds infiltration by cells was investigated using a confocal microscope (Zeiss LSM 880) and imaging in z-stack mode.

#### 4.6.3. Immunostaining and Fluorescence Microscopy

Immunostaining of the scaffolds for F-actin cytoskeleton, and αβ-crystallin, an HTM characteristic marker, was carried out on day 14. Pieces of the scaffolds were fixed in cold 4% PFA in PBS solution for 10 min, followed by 2-3 times rinsing in PBS for 5-10 min and stored at 4°C till staining.

Prior to staining, samples were permeabilizated with 0.5% Triton -X 100 in PBS for 15 min, and blocked with 0.1% Triton -X 100 in PBS mixed with 5% w/v BSA (denoted further as PBST solution) for 20 min.

Afterward, scaffolds were incubated in 1:200 Alexa fluor-488 Phalloidin (Thermofisher) for cytoskeleton staining and in 1:500 anti-αβ-crystallin (Abcam, Cambridge, MA) in PBST for 1 hour at room temperature. Samples were subsequently rinsed with PBST (three times) and incubated with secondary antibody Alexa flour-594 goat antimouse (Thermofisher, 1:100 dilution) for αβ-crystallin detection. Finally, samples were rinsed with PBST (2x), incubated with 1:1000 DAPI (Thermofisher) in PBS for 20 min for nuclei staining, and rinsed in PBS (2x). Imaging was performed using Nikon Ti-Eclipse (Nikon Instruments Europe B.V., Germany) with a Sola SE 365 II (Lumencor Inc., Beaverton, USA) solid-state illumination device and an Andor Clara CCD camera for detection.

#### 4.6.4. SEM of Cell Loaded Scaffolds

Similar imaging conditions as described above for scaffolds without cells were used. The scaffolds with cells were fixed using 2% glutaraldehyde in phosphate buffer, dehydrated, and dried using a graded series of increasing water/ethanol mixtures and hexamethyldisilazane (HDMS) before imaging. The scaffolds were incubated for 10 min in 50% v/v ethanol-water, 10 min in 70% v/v ethanol-water, 10 min in 80% v/v ethanol-water, 10 min in 90% v/v ethanol-water, 10 min in 96% v/v ethanol-water, 2×20 min in 100% ethanol, 10 min in 50% v/v ethanol-HMDS, 10 min in 100% HMDS and dried under ambient conditions overnight.

### 4.7. Statistical Analysis

All the results are reported as the mean ± standard deviation. Statistical differences were analyzed using one-way Analysis of Variance (ANOVA) and post-hoc Tuckey test, or unpaired t-test, performed with In Stat3 software. Differences with p < 0.05 were considered significant.

## Acknowledgements

The authors thank Spoorti Ramesh from the University of Saarland and INM Leibniz Institute for New Materials, Saarbrücken, Germany, Lizbeth Ofelia Prieto-López, Robert Drumm, Marlon Jochum and Ha Rimbach from INM Leibniz Institute for New Materials, Saarbrücken, Germany for the support in measurements and data collection, and Lukas Schwab from TA Instruments for the support in rheological data analysis. MV and AdC acknowledge financial support from the EU within BioSmartTrainee, the Marie Skłodowska-Curie Innovative Training School, No. 642861. PW acknowledges the China Scholarship Council (CSC).

---

Human Trabecular Meshwork (HTM) is a gradient structure located in the eye, responsible for maintaining proper pressure inside the eyeball. HTM dysfunctions lead to glaucoma. By employing melt electrowriting (MEW, we aim at the reconstruction of the complexity of the native HTM tissue. The present work demonstrates the utility of MEW to reconstruct complex morphological features of natural tissues.

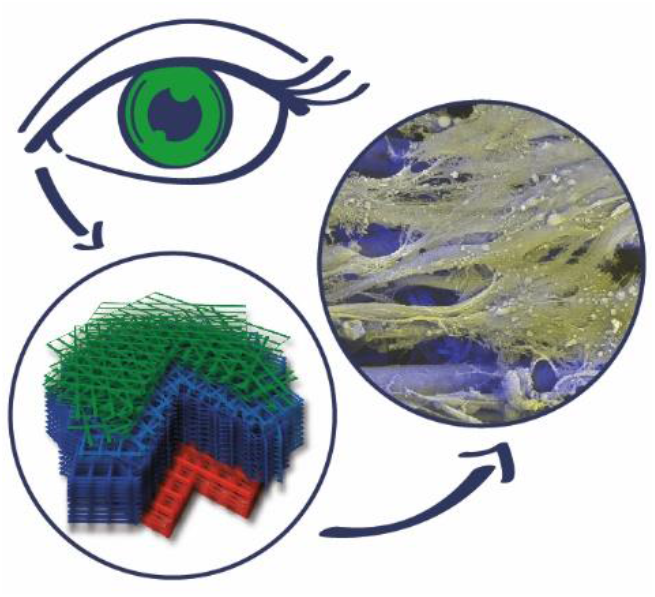

## References

[1] A. Seidi, M. Ramalingam, I. Elloumi-Hannachi, S. Ostrovidov, A. Khademhosseini, Gradient biomaterials for soft-to-hard interface tissue engineering, Acta Biomaterialia 7(4) (2011) 1441–1451.

[2] R. Obregón, J. Ramón-Azcón, S. Ahadian, H. Shiku, M. Ramalingam, A. Khademhosseini, T. Matsue, Chapter 13 - Gradient Biomaterials as Tissue Scaffolds, in: A. Vishwakarma, P. Sharpe, S. Shi, M. Ramalingam (Eds.), Stem Cell Biology and Tissue Engineering in Dental Sciences, Academic Press, Boston, 2015, pp. 175–186.

[3] W.D. Stamer, A.F. Clark, The many faces of the trabecular meshwork cell, Experimental eye research 158 (2017) 112–123.

[4] J.C.H. Tan, J.M. Gonzalez, Jr., S. Hamm-Alvarez, J. Song, In Situ Autofluorescence Visualization of Human Trabecular Meshwork Structure, Investigative Ophthalmology & Visual Science 53(4) (2012) 2080–2088.

[5] D.W. Abu-Hassan, T.S. Acott, M.J. Kelley, The Trabecular Meshwork: A Basic Review of Form and Function, Journal of ocular biology 2(1) (2014).

[6] W.B. Trattler, P.K. Kaiser, N.J. Friedman, Review of ophthalmology E-book: Expert consult-online and print, Elsevier Health Sciences 2012.

[7] C.N. Dautriche, Y. Xie, S.T. Sharfstein, Walking through trabecular meshwork biology: Toward engineering design of outflow physiology, Biotechnology advances 32(5) (2014) 971–83.

[8] K.Y. Torrejon, D. Pu, M. Bergkvist, J. Danias, S.T. Sharfstein, Y. Xie, Recreating a human trabecular meshwork outflow system on microfabricated porous structures, Biotechnology and bioengineering 110(12) (2013) 3205–18.

[9] T.S. Dietlein, P.C. Jacobi, C. Lüke, G.K. Krieglstein, Morphological variability of the trabecular meshwork in glaucoma patients: implications for non-perforating glaucoma surgery, The British journal of ophthalmology 84(12) (2000) 1354–9.

[10] Y. Sundaresan, M. Veerappan, K.S. Ramasamy, G.P. Chidambaranathan, Identification, quantification and age-related changes of human trabecular meshwork stem cells, Eye and vision (London, England) 6 (2019) 31.

[11] A.M. Demetriades, Gene therapy for glaucoma, Current opinion in ophthalmology 22(2) (2011) 73–7.

[12] J.A. Last, T. Pan, Y. Ding, C.M. Reilly, K. Keller, T.S. Acott, M.P. Fautsch, C.J. Murphy, P. Russell, Elastic modulus determination of normal and glaucomatous human trabecular meshwork, Investigative ophthalmology & visual science 52(5) (2011) 2147–2152.

[13] H. Li, T. Bagué, A. Kirschner, A.N. Strat, H. Roberts, R.W. Weisenthal, A.E. Patteson, N. Annabi, W.D. Stamer, P.S. Ganapathy, S. Herberg, A tissue-engineered human trabecular meshwork hydrogel for advanced glaucoma disease modeling, Experimental eye research 205 (2021) 108472.

[14] K. Wang, M.A. Johnstone, C. Xin, S. Song, S. Padilla, J.A. Vranka, T.S. Acott, K. Zhou, S.A. Schwaner, R.K. Wang, T. Sulchek, C.R. Ethier, Estimating Human Trabecular Meshwork Stiffness by Numerical Modeling and Advanced OCT Imaging, Invest Ophthalmol Vis Sci 58(11) (2017) 4809–4817.

[15] K. Wang, A.T. Read, T. Sulchek, C.R. Ethier, Trabecular meshwork stiffness in glaucoma, Experimental eye research 158 (2017) 3–12.

[16] B. Dhandayuthapani, Y. Yoshida, T. Maekawa, D.S. Kumar, Polymeric Scaffolds in Tissue Engineering Application: A Review, International Journal of Polymer Science 2011 (2011) 290602.

[17] O.E. Armitage, M.L. Oyen, Hard-Soft Tissue Interface Engineering, in: L.E. Bertassoni, P.G. Coelho (Eds.), Engineering Mineralized and Load Bearing Tissues, Springer International Publishing, Cham, 2015, pp. 187–204.

[18] K.Y. Torrejon, E.L. Papke, J.R. Halman, M. Bergkvist, J. Danias, S.T. Sharfstein, Y. Xie, TGFβ2-induced outflow alterations in a bioengineered trabecular meshwork are offset by a rho-associated kinase inhibitor, Scientific Reports 6(1) (2016) 38319.

[19] K.Y. Torrejon, E.L. Papke, J.R. Halman, J. Stolwijk, C.N. Dautriche, M. Bergkvist, J. Danias, S.T. Sharfstein, Y. Xie, Bioengineered glaucomatous 3D human trabecular meshwork as an in vitro disease model, Biotechnology and bioengineering 113(6) (2016) 1357–68.

[20] M. Osmond, S.M. Bernier, M.B. Pantcheva, M.D. Krebs, Collagen and collagen-chondroitin sulfate scaffolds with uniaxially aligned pores for the biomimetic, three dimensional culture of trabecular meshwork cells, Biotechnology and bioengineering 114(4) (2017) 915–923.

[21] M.J. Osmond, M.D. Krebs, M.B. Pantcheva, Human trabecular meshwork cell behavior is influenced by collagen scaffold pore architecture and glycosaminoglycan composition, Biotechnology and bioengineering 117(10) (2020) 3150–3159.

[22] S.C. Saccà, S. Tirendi, S. Scarfì, M. Passalacqua, F. Oddone, C.E. Traverso, S. Vernazza, A.M. Bassi, An advanced in vitro model to assess glaucoma onset, Altex 37(2) (2020) 265–274.

[23] M. Bouchemi, C. Roubeix, K. Kessal, L. Riancho, A.L. Raveu, H. Soualmia, C. Baudouin, F. Brignole-Baudouin, Effect of benzalkonium chloride on trabecular meshwork cells in a new in vitro 3D trabecular meshwork model for glaucoma, Toxicology in vitro : an international journal published in association with BIBRA 41 (2017) 21–29.

[24] J. Buffault, A. Labbé, P. Hamard, F. Brignole-Baudouin, C. Baudouin, The trabecular meshwork: Structure, function and clinical implications. A review of the literature, Journal francais d’ophtalmologie 43(7) (2020) e217–e230.

[25] L. Baldino, S. Cardea, N. Maffulli, E. Reverchon, Regeneration techniques for bone-to-tendon and muscle-to-tendon interfaces reconstruction, British Medical Bulletin 117(1) (2016) 25–37.

[26] J. Rnjak-Kovacina, A.S. Weiss, Increasing the pore size of electrospun scaffolds, Tissue engineering. Part B, Reviews 17(5) (2011) 365–72.

[27] H. Chen, A.d.B.F.B. Malheiro, C. van Blitterswijk, C. Mota, P.A. Wieringa, L. Moroni, Direct Writing Electrospinning of Scaffolds with Multidimensional Fiber Architecture for Hierarchical Tissue Engineering, ACS Applied Materials & Interfaces 9(44) (2017) 38187–38200.

[28] P.D. Dalton, Melt electrowriting with additive manufacturing principles, Current Opinion in Biomedical Engineering 2 (2017) 49–57.

[29] T.D. Brown, P.D. Dalton, D.W. Hutmacher, Direct Writing By Way of Melt Electrospinning, Advanced Materials 23(47) (2011) 5651–5657.

[30] A. Youssef, A. Hrynevich, L. Fladeland, A. Balles, J. Groll, P.D. Dalton, S. Zabler, The Impact of Melt Electrowritten Scaffold Design on Porosity Determined by X-Ray Microtomography, Tissue Engineering Part C: Methods 25(6) (2019) 367–379.

[31] A. Hrynevich, B. Elçi, J.N. Haigh, R. McMaster, A. Youssef, C. Blum, T. Blunk, G. Hochleitner, J. Groll, P.D. Dalton, Dimension-Based Design of Melt Electrowritten Scaffolds, Small 14(22) (2018) e1800232.

[32] C. Xie, Q. Gao, P. Wang, L. Shao, H. Yuan, J. Fu, W. Chen, Y. He, Tendril Climber Inspired Structure-Induced Cell Growth by Direct Writing Heterogeneous Scaffold, Available at SSRN 3321957 (2019).

[33] M. Castilho, D. Feyen, M. Flandes-Iparraguirre, G. Hochleitner, J. Groll, P.A.F. Doevendans, T. Vermonden, K. Ito, J.P.G. Sluijter, J. Malda, Melt Electrospinning Writing of Poly-Hydroxymethylglycolide-co-ε-Caprolactone-Based Scaffolds for Cardiac Tissue Engineering, Adv Healthc Mater 6(18) (2017).

[34] K.F. Eichholz, D.A. Hoey, Mediating human stem cell behaviour via defined fibrous architectures by melt electrospinning writing, Acta Biomaterialia 75 (2018) 140–151.

[35] M. Gwiazda, S. Kumar, W. Świeszkowski, S. Ivanovski, C. Vaquette, The effect of melt electrospun writing fiber orientation onto cellular organization and mechanical properties for application in Anterior Cruciate Ligament tissue engineering, Journal of the Mechanical Behavior of Biomedical Materials 104 (2020) 103631.

[36] N.T. Saidy, F. Wolf, O. Bas, H. Keijdener, D.W. Hutmacher, P. Mela, E.M. De-Juan-Pardo, Biologically Inspired Scaffolds for Heart Valve Tissue Engineering via Melt Electrowriting, Small 15(24) (2019) 1900873.

[37] N.T. Nguyen, J.H. Kim, Y.H. Jeong, Identification of sagging in melt-electrospinning of microfiber scaffolds, Materials science & engineering. C, Materials for biological applications 103 (2019) 109785.

[38] N. Abbasi, A. Abdal-hay, S. Hamlet, E. Graham, S. Ivanovski, Effects of Gradient and Offset Architectures on the Mechanical and Biological Properties of 3-D Melt Electrowritten (MEW) Scaffolds, ACS Biomaterials Science & Engineering 5(7) (2019) 3448–3461.

[39] Y. Han, M. Lian, B. Sun, B. Jia, Q. Wu, Z. Qiao, K. Dai, Preparation of high precision multilayer scaffolds based on Melt Electro-Writing to repair cartilage injury, Theranostics 10(22) (2020) 10214–10230.

[40] Y. Han, B. Jia, M. Lian, B. Sun, Q. Wu, B. Sun, Z. Qiao, K. Dai, High-precision, gelatin-based, hybrid, bilayer scaffolds using melt electro-writing to repair cartilage injury, Bioactive Materials 6(7) (2021) 2173–2186.

[41] E. Hewitt, S. Mros, M. McConnell, J.D. Cabral, A. Ali, Melt-electrowriting with novel milk protein/PCL biomaterials for skin regeneration, Biomedical materials (Bristol, England) 14(5) (2019) 055013.

[42] M. von Witzleben, T. Stoppe, T. Ahlfeld, A. Bernhardt, M.L. Polk, M. Bornitz, M. Neudert, M. Gelinsky, Biomimetic Tympanic Membrane Replacement Made by Melt Electrowriting, Advanced Healthcare Materials 10(10) (2021).

[43] G. Hochleitner, E. Fürsattel, R. Giesa, J. Groll, H.-W. Schmidt, P.D. Dalton, Melt Electrowriting of Thermoplastic Elastomers, Macromolecular Rapid Communications 39(10) (2018) 1800055.

[44] J.C. Kade, P.D. Dalton, Polymers for Melt Electrowriting, Advanced Healthcare Materials 10(1) (2021) 2001232.

[45] H. Sun, L. Mei, C. Song, X. Cui, P. Wang, The in vivo degradation, absorption and excretion of PCL-based implant, Biomaterials 27(9) (2006) 1735–40.

[46] G. Collins, J. Federici, Y. Imura, L.H. Catalani, Charge generation, charge transport, and residual charge in the electrospinning of polymers: A review of issues and complications, Journal of Applied Physics 111(4) (2012) 044701.

[47] C. Vaquette, J.J. Cooper-White, Increasing electrospun scaffold pore size with tailored collectors for improved cell penetration, Acta Biomaterialia 7(6) (2011) 2544–2557.

[48] C. Vaquette, J. Cooper-White, A simple method for fabricating 3-D multilayered composite scaffolds, Acta biomaterialia 9(1) (2013) 4599–4608.

[49] D.W. Hutmacher, Scaffolds in tissue engineering bone and cartilage, Biomaterials 21(24) (2000) 2529–2543.

[50] C.-C. Yeh, C.-N. Chen, Y.-T. Li, C.-W. Chang, M.-Y. Cheng, H.-I. Chang, The Effect of Polymer Molecular Weight and UV Radiation on Physical Properties and Bioactivities of PCL Films, Cellular Polymers 30(5) (2011) 261–276.

[51] Y.A. Mehanna, E. Sadler, R.L. Upton, A.G. Kempchinsky, Y. Lu, C.R. Crick, The challenges, achievements and applications of submersible superhydrophobic materials, Chemical Society Reviews 50(11) (2021) 6569–6612.

[52] M.A. Woodruff, D.W. Hutmacher, The return of a forgotten polymer—Polycaprolactone in the 21st century, Progress in Polymer Science 35(10) (2010) 1217–1256.

[53] Y. Rotbaum, C. Puiu, D. Rittel, M. Domingos, Quasi-static and dynamic in vitro mechanical response of 3D printed scaffolds with tailored pore size and architectures, Materials Science and Engineering: C 96 (2019) 176–182.

[54] M.M. Mirhosseini, V. Haddadi-Asl, S.S. Zargarian, Fabrication and characterization of hydrophilic poly(ε-caprolactone)/pluronic P123 electrospun fibers, Journal of Applied Polymer Science 133(17) (2016).

[55] M. Castilho, G. Hochleitner, W. Wilson, B. van Rietbergen, P.D. Dalton, J. Groll, J. Malda, K. Ito, Mechanical behavior of a soft hydrogel reinforced with three-dimensional printed microfibre scaffolds, Scientific Reports 8(1) (2018) 1245.

[56] L.J. Camras, W.D. Stamer, D. Epstein, P. Gonzalez, F. Yuan, Differential Effects of Trabecular Meshwork Stiffness on Outflow Facility in Normal Human and Porcine Eyes, Investigative Ophthalmology & Visual Science 53(9) (2012) 5242–5250.

[57] L.J. Camras, W.D. Stamer, D. Epstein, P. Gonzalez, F. Yuan, Circumferential Tensile Stiffness of Glaucomatous Trabecular Meshwork, Investigative Ophthalmology & Visual Science 55(2) (2014) 814–823.

[58] D.W. Hutmacher, T. Schantz, I. Zein, K.W. Ng, S.H. Teoh, K.C. Tan, Mechanical properties and cell cultural response of polycaprolactone scaffolds designed and fabricated via fused deposition modeling, Journal of Biomedical Materials Research 55(2) (2001) 203–216.

