## Supplementary Information for "Melt Electrowriting of Scaffolds with a Porosity Gradient to Mimic the Matrix Structure of the Human Trabecular Meshwork"

**Table S1.** Molecular weight and thermal properties of PCL before processing, after melting (and subsequent cooling, but no printing), and printing.

| Sample | Tm ($^{oC}$) | Crystallinity fraction (%) | *Mn* | *Mw* | Polydispersity (*Mw/Mn)* |
|---|---|---|---|---|---|
| PCL raw | 57.1 | 84.6 | 55475 | 102000 | 1.8387 |
| PCL after melting | 57.5 | 78.8 | 68596 | 101760 | 1.4835 |
| PCL after printing | 58.3 | 87.4 | 66934 | 108180 | 1.6162 |

crystallinity was calculated from the enthalpy value of the melting peak, assuming 139.5 J/g for 100% crystallinity (based on [1])

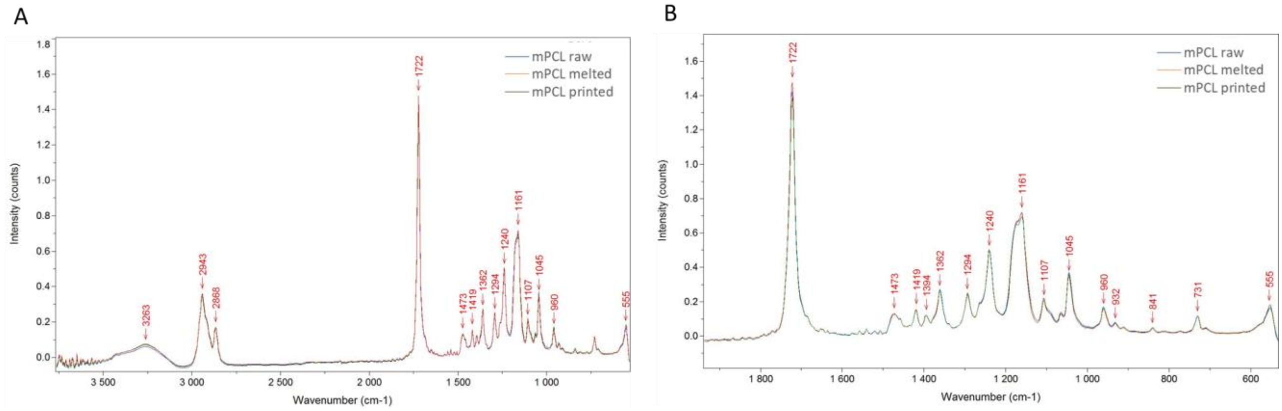


**Figure S1.** FTIR spectra of material in a raw state, after melting, and after printing.

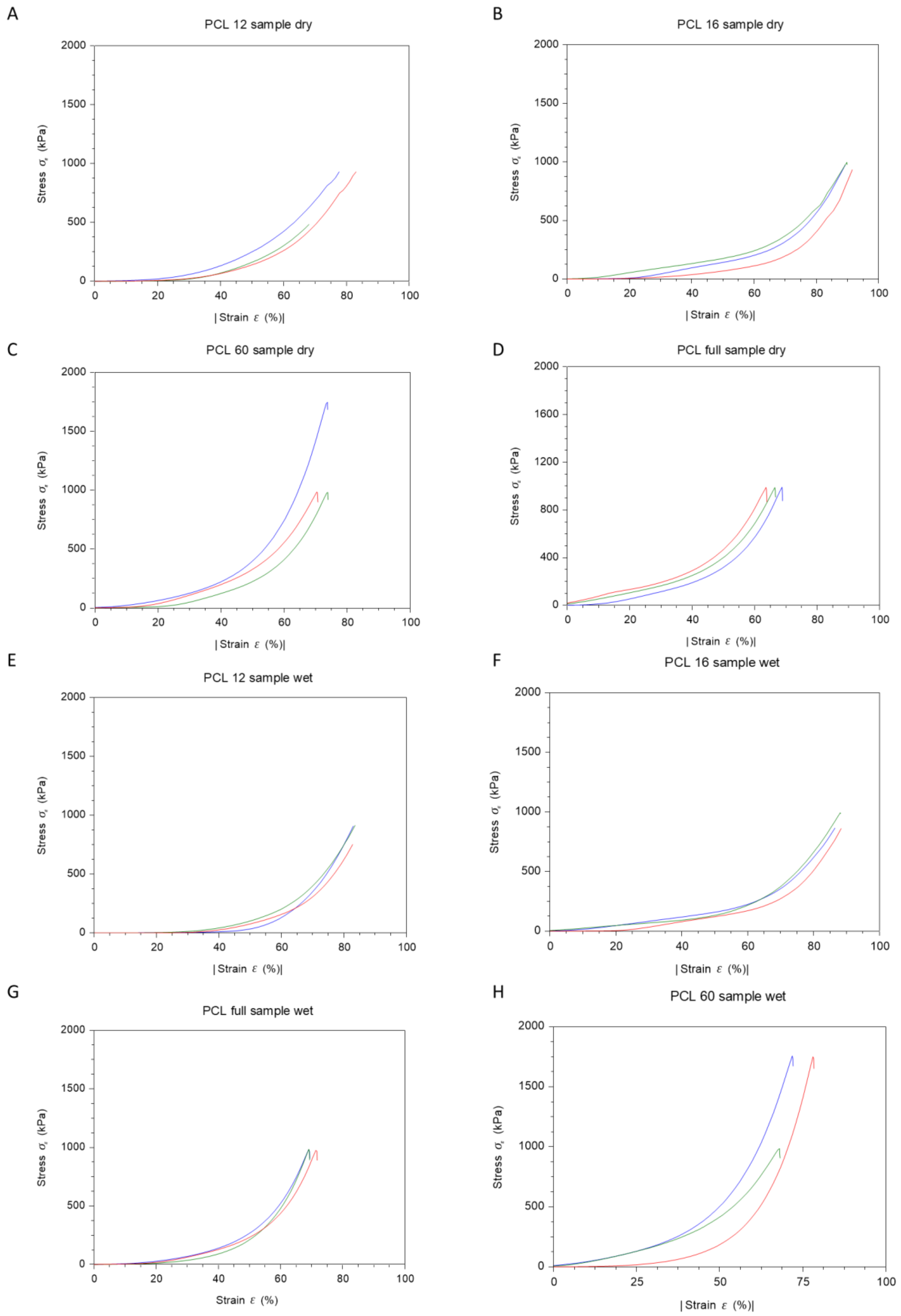


**Figure S2.** The results of the compression test for all measured samples in the dry state: PCL 12 (A), PCL 16 (B), PCL 60 (C), PCL full (D), and wet state PCL 12 (E), PCL 16 (F), PCL 60 (G), PCL full (H), to present samples diversity.

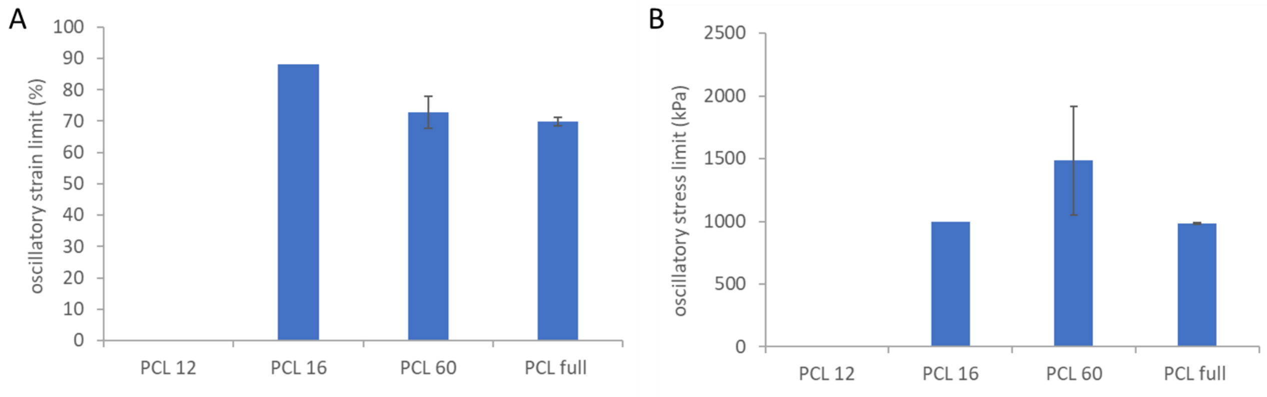


**Figure S3.** Compression testing of different scaffolds. Oscillatory stress (A) and strain (B) limits for different samples. For PCL 12, the limit was not detected, for PCL 16 only for one sample, which we assigned to the small sample thickness.

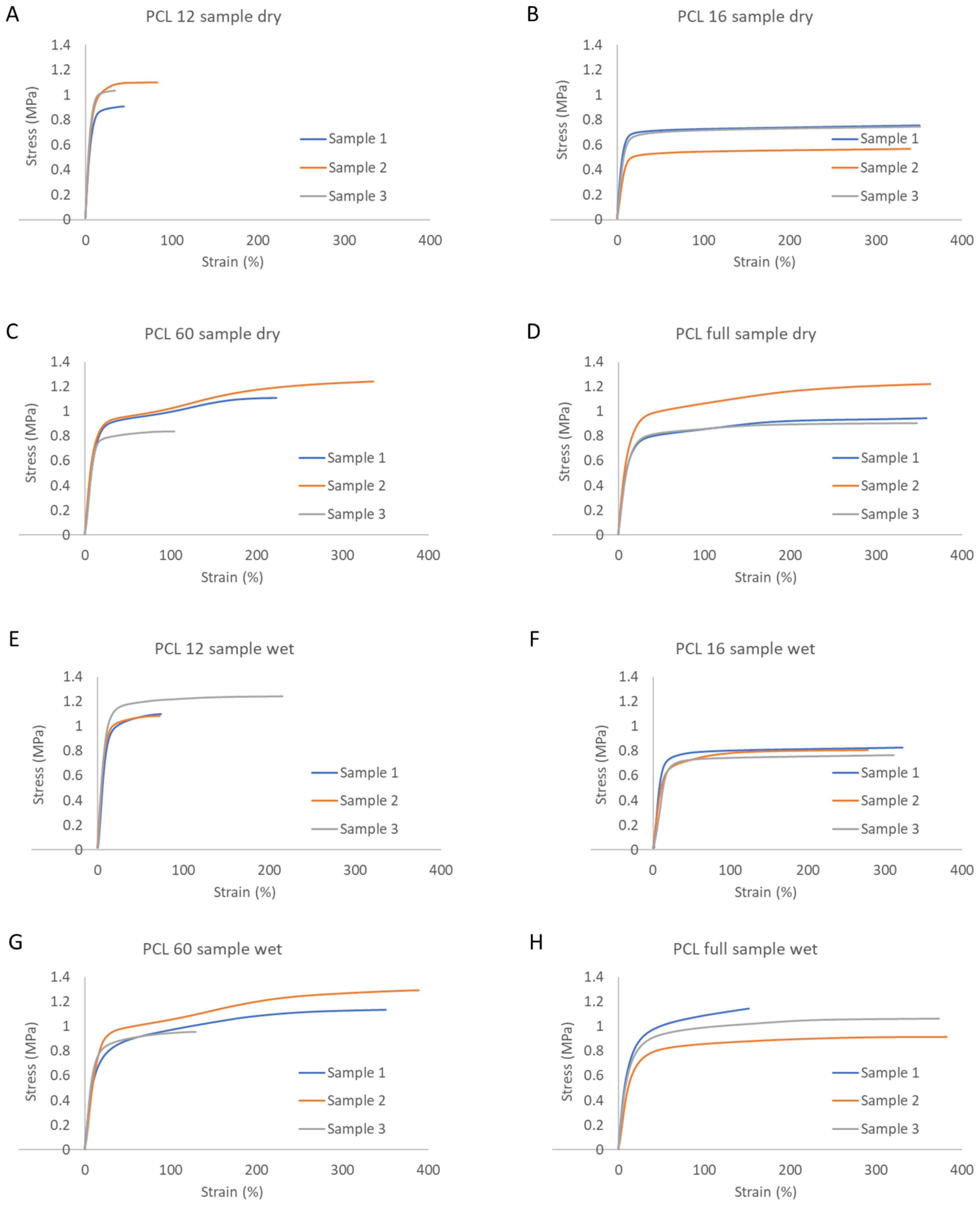


**Figure S4.** The results of the tensile test for all measured samples in the dry state: PCL 12 (A), PCL 16 (B), PCL 60 (C), PCL full (D), and wet state PCL 12 (E), PCL 16 (F), PCL 60 (G), PCL full (H), to present samples diversity.

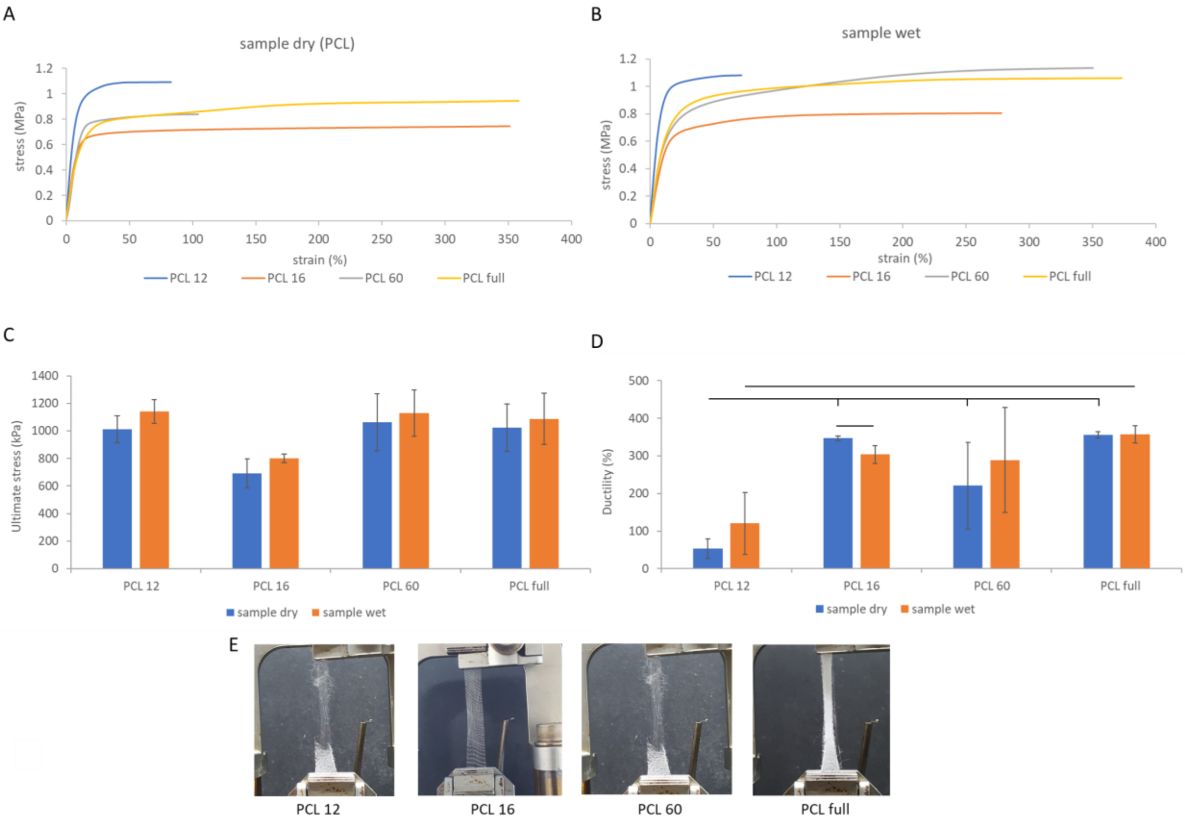


**Figure S5.** Tensile testing of different scaffolds. Full stress-strain curves of the scaffolds tested in a dry state (A), and a wet state (B). Comparison of ultimate stress (C) and ductility (D). Typical samples at the end of the test, after ca. 300% elongation (E).

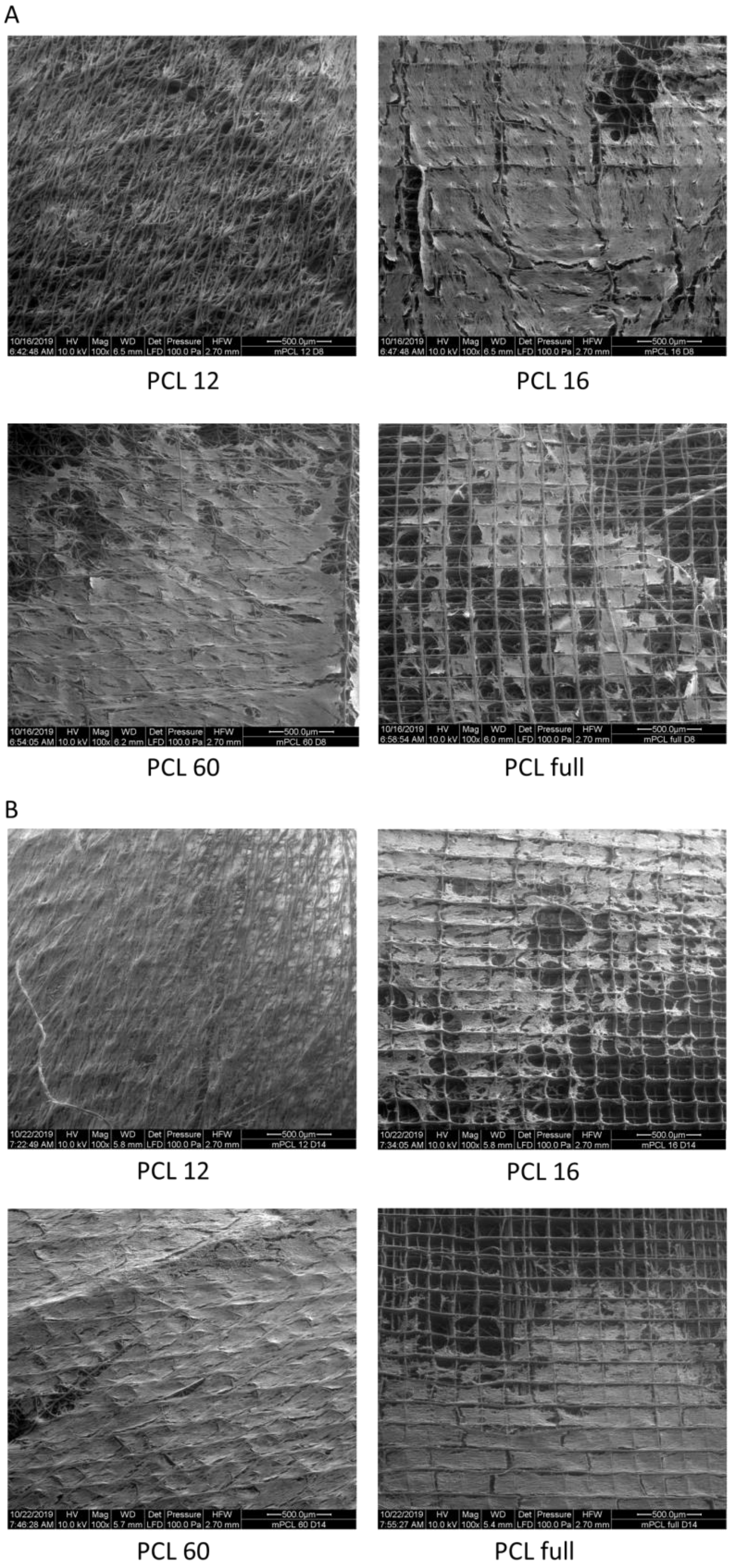


**Figure S6.** SEM images of the PCL scaffolds at mag. 100x after 8 days (A) and 14 days (B) of cell culture (presented at higher magnification in Figure 6 in the main text of the manuscript). The ability of the cells to migrate along the scaffolds is revealed.

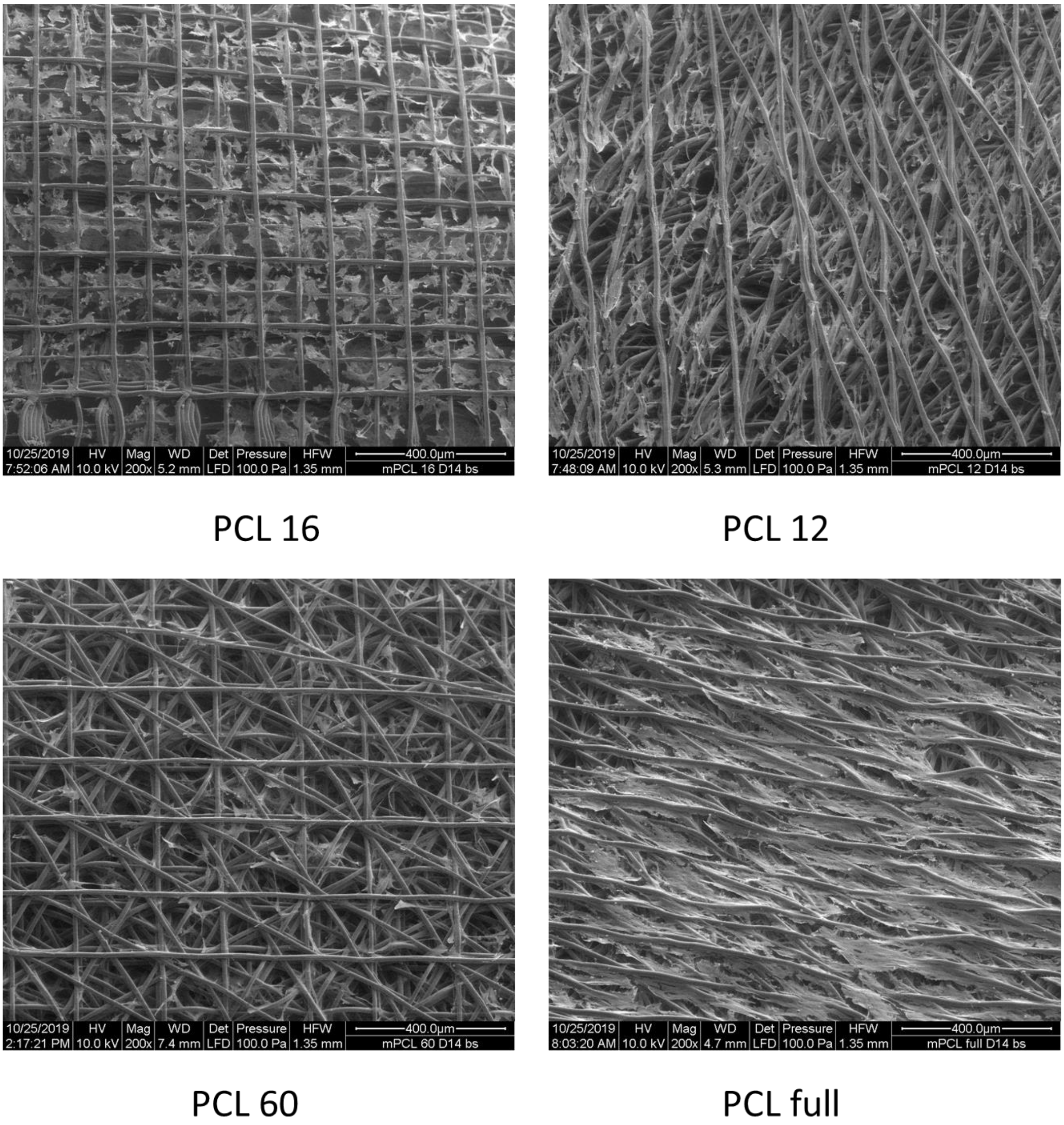


**Figure S7.** SEM images of the PCL scaffolds at mag. 200x after 14 days of cell culture. Scaffolds were imaged from the bottom side with respect to the position during cell experiments and the reverse side compared to Figure 5 in the main text of the manuscript. The ability of the cells to penetrate throughout the scaffolds is revealed.

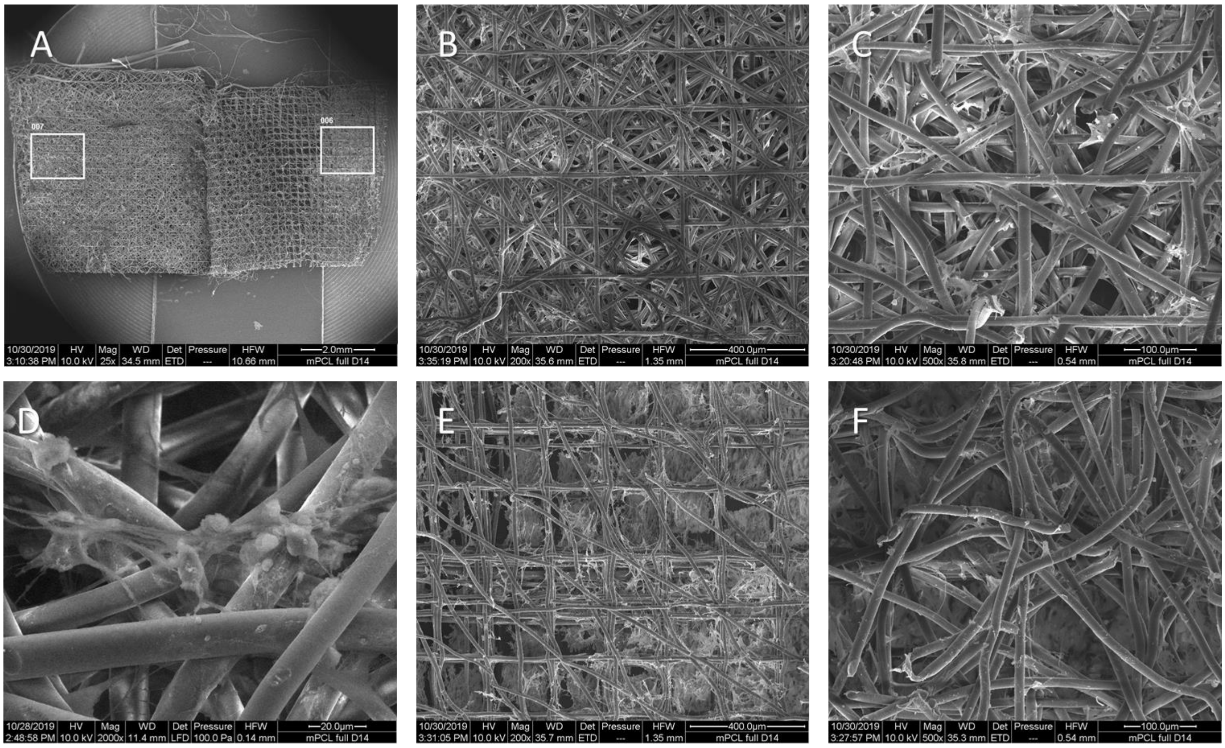


**Figure S8.** SEM images on the PCL full scaffolds, after horizontal cut through the middle of the structure. Scaffold after cutting, with the bottom part on the left and the top part on the right, and marked regions of further imaging (A). Interior of the scaffold with visible cells: bottom side (B-D), top side (E-F). The ability of the cells to penetrate throughout the scaffold and habituate the middle phase is revealed.

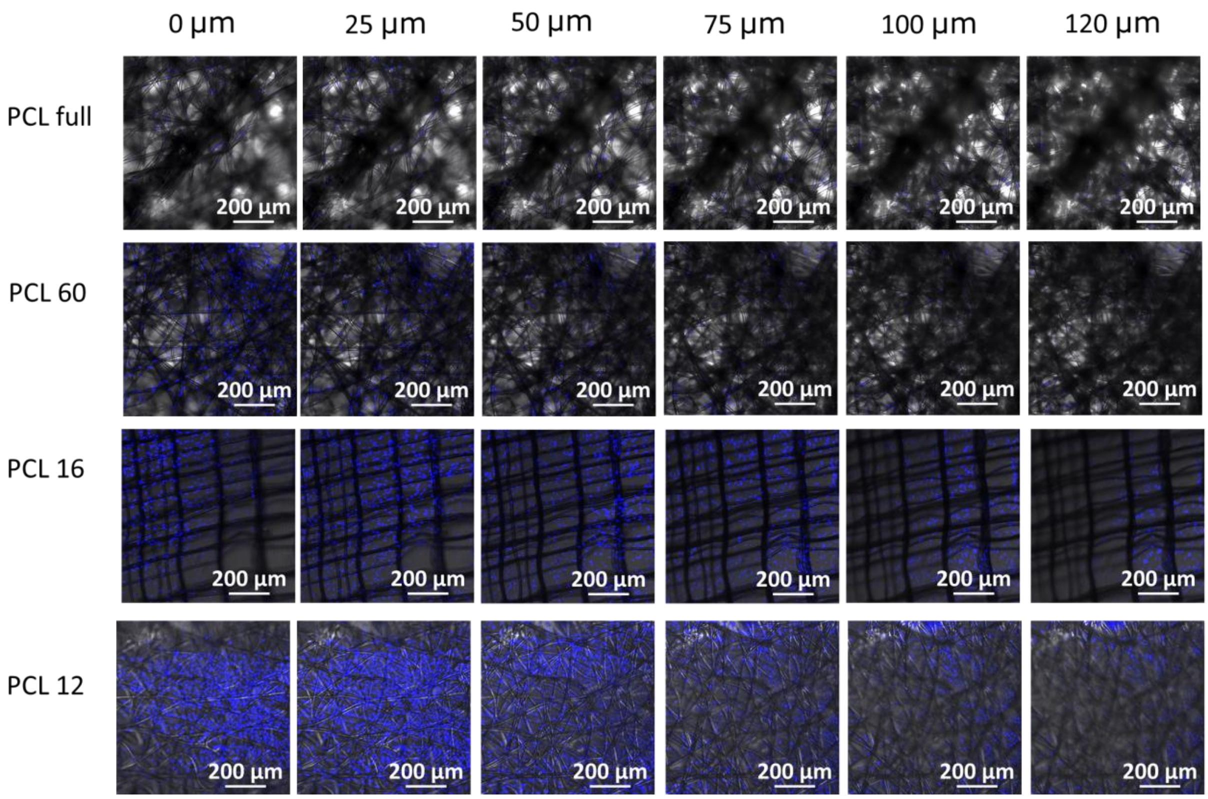


**Figure S9.** Confocal images of the scaffolds seeded with cells, after 14 days of cell culture, at different z-levels. In blue, the nuclei stained with DAPI are visible. Smaller cell density observed in PCL full scaffolds can be connected to the difficulty in light microscopic observation in the thicker, non-transparent scaffolds.

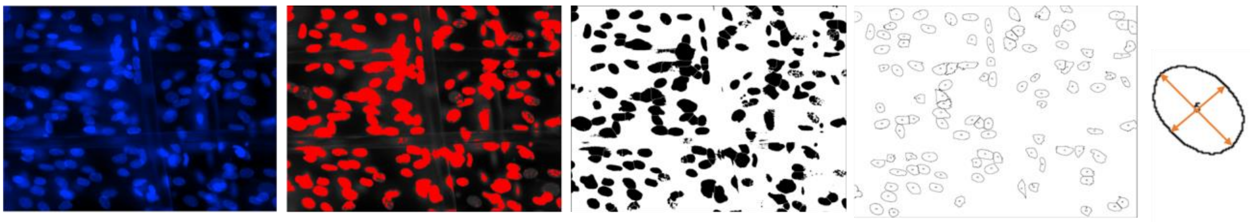

**Figure S10.** Exemplary image processing used for the nuclei aspect ratio quantification: original image, image after adjustment of the threshold, image after applying “watershed” function on a binary copy, nuclei considered in the calculation after applying size exclusion limit (50 - 150 μm), width and length of the nuclei. Note that the nuclei in case of clamps of cells or overlapping cells were excluded from the calculation.

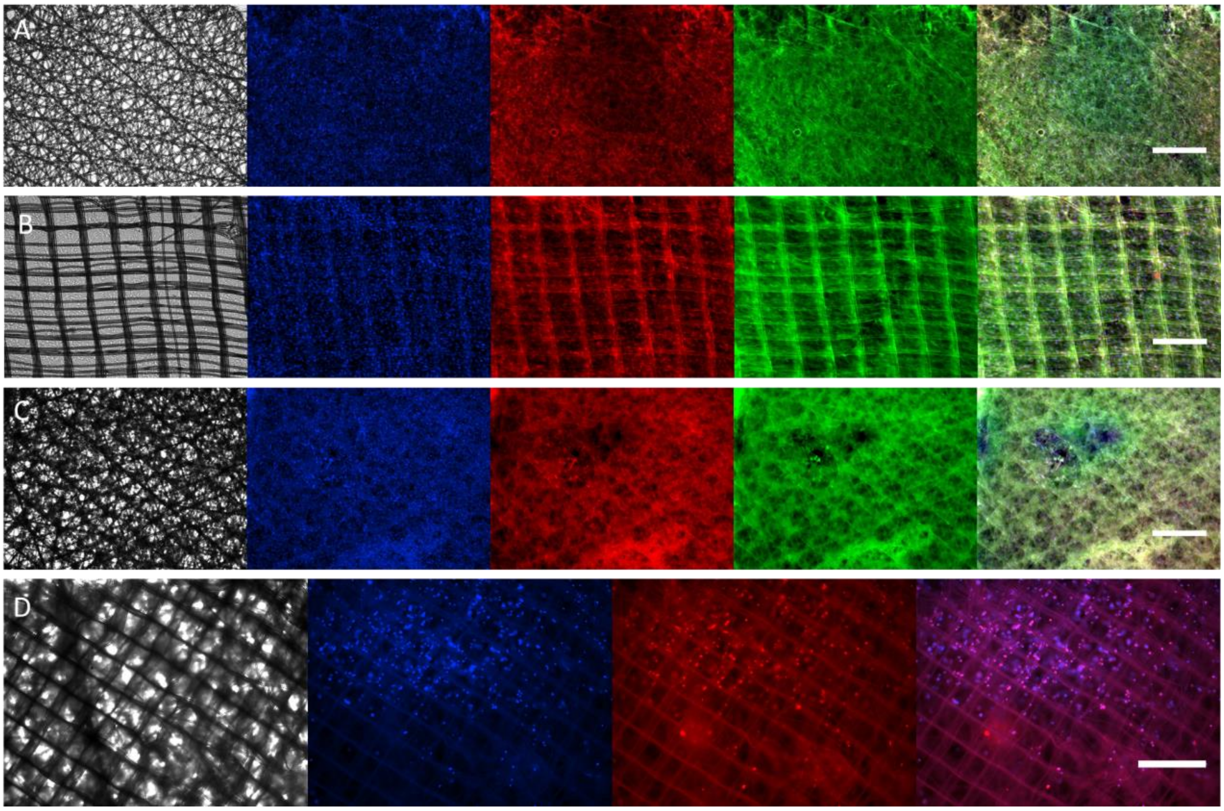


**Figure S11.** Fluorescent images of the cells cultured on different scaffold types at lower magnification (4x): PCL 12 (A), PCL 16 (B), PCL 60 (C), PCL full (D). The gray channel represents a bright field, the blue channel DAPI nuclei staining, the red channel ab-crystallin, the green channel phalloidin actin staining, and the last tile is a merged representation. Scale bars 500 µm. The difficulty of light microscopy imaging of the thicker scaffolds (D) is caused by the light reflection on the thicker, non-transparent designs.

**References:**

[1] D.W. Hutmacher, T. Schantz, I. Zein, K.W. Ng, S.H. Teoh, K.C. Tan, Mechanical properties and cell cultural response of polycaprolactone scaffolds designed and fabricated via fused deposition modeling, Journal of Biomedical Materials Research 55(2) (2001) 203-216.
